# Liquid condensate is a common state of proteins and polypeptides at the regime of high intermolecular interactions

**DOI:** 10.1101/2021.12.31.474648

**Authors:** Manisha Poudyal, Komal Patel, Ajay Singh Sawner, Laxmikant Gadhe, Pradeep Kadu, Debalina Datta, Semanti Mukherjee, Soumik Ray, Ambuja Navalkar, Siddhartha Maiti, Debdeep Chatterjee, Riya Bera, Nitisha Gahlot, Ranjith Padinhateeri, Samir K. Maji

## Abstract

Liquid-liquid phase separation (LLPS) has emerged as a crucial biological mechanism for sequestering macromolecules (such as proteins and nucleic acids) into membraneless organelles in cells. Unstructured and intrinsically disordered domains are known to facilitate multivalent interactions driving protein LLPS. We hypothesized that LLPS could be an intrinsic property of proteins/polypeptides at their high intermolecular interaction regime. To examine this, we studied many (a total of 23) proteins/polypeptides with different structures and sequences for LLPS study using molecular crowder polyethylene glycol (PEG-8000). We showed that all proteins and even highly charged polypeptides (under study) can undergo liquid condensate formation, however with different phase space and conditions. Using a single component and combinations of protein multicomponent (co-LLPS) systems, we establish that a variety of intermolecular interactions can drive proteins/polypeptides LLPS.

## Introduction

Liquid-liquid phase separation (LLPS) of biomolecules (proteins/nucleic acids) is now well-established as a ubiquitous mechanism for the formation of membraneless organelles^1-6^. These phase separated, condensed compartments not only help in various cellular functionality^7-9^; but they are also useful for macromolecular sequestration/storage, and cellular signaling/communications^1,9^. Although LLPS substantially contributes towards the overall fitness of cells^1,2,10,11^; it is also associated with certain risks^1,6,12,13^. Concentrations of protein increase several orders of magnitude inside the condensate^14-16^ compared to their endogenous levels. This often leads to toxic protein aggregation and nucleation of amyloid fibril formation associated with various human diseases such as amyotrophic lateral sclerosis (ALS), Alzheimer’s disease (AD), and Parkinson’s disease (PD)^1,17-22^. It is widely accepted that intra- and inter-molecular interactions driving protein phase separation are embedded in the protein/peptide sequence and the respective structure ^1,4,6,14,23^. In this context, the conformational properties of intrinsically disordered regions (IDRs), low complexity domains (LCDs), and prion-like domains (PLDs) are known to facilitate multivalent interactions that are prerequisites for condensate formation^23-28^. Further, the specific arrangement of amino acids in protein sequences (e.g., spacers and stickers) under various conditions can regulate LLPS^23,27,29^ through common mechanisms that promote these multivalent interactions (such as electrostatic and cation-π interactions)^23,25,26,29,30^. The nature of the overall interactions between the proteins makes the condensates responsive to the cellular microenvironment^31,32^. By exploiting this knowledge, it is also possible to design artificial peptides/proteins with tunable LLPS properties^8,33,34^.

However, emerging evidence indicates that a significant proportion of proteins in the human proteome reside at concentrations just below their respective solubility limit^35^. The concentration levels not only depend on the extent of endogenous expression of individual proteins; but can also be greatly affected by the efficiency of the protein turnover machinery of the cell. The transition from soluble to LLPS state (reaching the critical concentration) thus, is not associated with a very high energy barrier^2,32,36,37^. Seemingly, alterations such as post-translational modifications, changes in cellular or subcellular localization, the effect of counterions, and metabolites (such as ATP) can significantly modulate the phase behavior of various proteins^5,30,38-42^. Apart from intrinsically disordered proteins (IDPs), globular proteins (such as lysozyme^43^) are also capable of undergoing LLPS. Since the basis of most supramolecular assemblies (aggregates/precipitates/LLPS/crystals) is the intermolecular interactions, by tuning the extent of such interactions, it is experimentally feasible to explore conditions that drive LLPS of globular proteins as well. Based on this, we hypothesize that LLPS could be a generic state of proteins/polypeptides when a threshold intermolecular interaction is achieved. However, their relative extent of LLPS might differ due to the overall intermolecular interaction strength.

To prove our hypothesis, we chose 19 different proteins with diverse structures/sequences and subjected them to macromolecular crowding using PEG-8000. PEG is widely used as an inducer for protein LLPS due to its inert nature and provides a crowding effect similar to cellular milieu. We observed that all the proteins under study could undergo thermo-reversible LLPS with different critical concentrations and also with different kinetics. The relative extent of LLPS depends on their intermolecular interaction strength as well as solubility. Further, our data suggests that the proteins might use one or a combination of the interaction modes (electrostatic/hydrophobic/H-bonding) to form phase separated condensates. We further designed polypeptides based on neutral (Gly), hydrophobic (Val), positively charged (Arg), and negatively charged (Asp) amino acids and observed that they also undergo phase separation at conditions favorable for their intermolecular interactions. Moreover, the heterotypic LLPS (co-LLPS) by various combination of proteins suggests that LLPS/co-LLPS might be the result of intermolecular interactions by proteins/polypeptides, which might be controlled by various intrinsic and extrinsic factors including protein concentration. The present study indicates that LLPS might be a shared property of proteins/polypeptides. However, the soluble versus LLPS state could be tightly regulated by cells depending on the cellular requirement^6,44-48^.

## Results

### Liquid-liquid phase separation (LLPS) of a diverse library of proteins

To address if LLPS is a generic phenomenon of proteins, we first examined whether a diverse library of proteins could undergo LLPS *in vitro* in the presence of a molecular crowder (in our case, PEG-8000). We chose this library of proteins from multiple species with varied sequences, structures and properties (Table S1). Also, to exclude the influence of various cellular factors and other parameters such as salt, we have used 20 mM sodium phosphate buffer (pH 7.4) in the presence of PEG-8000 as a molecular crowder. We first generated the three-dimensional surface image of proteins superimposed with their secondary structures using PYMOL (v 2.5.2) to understand the diversity of structures and distribution of charge (Fig. 1a and S1). We then examined all the protein sequences *in silico*, using

**Figure 1.**
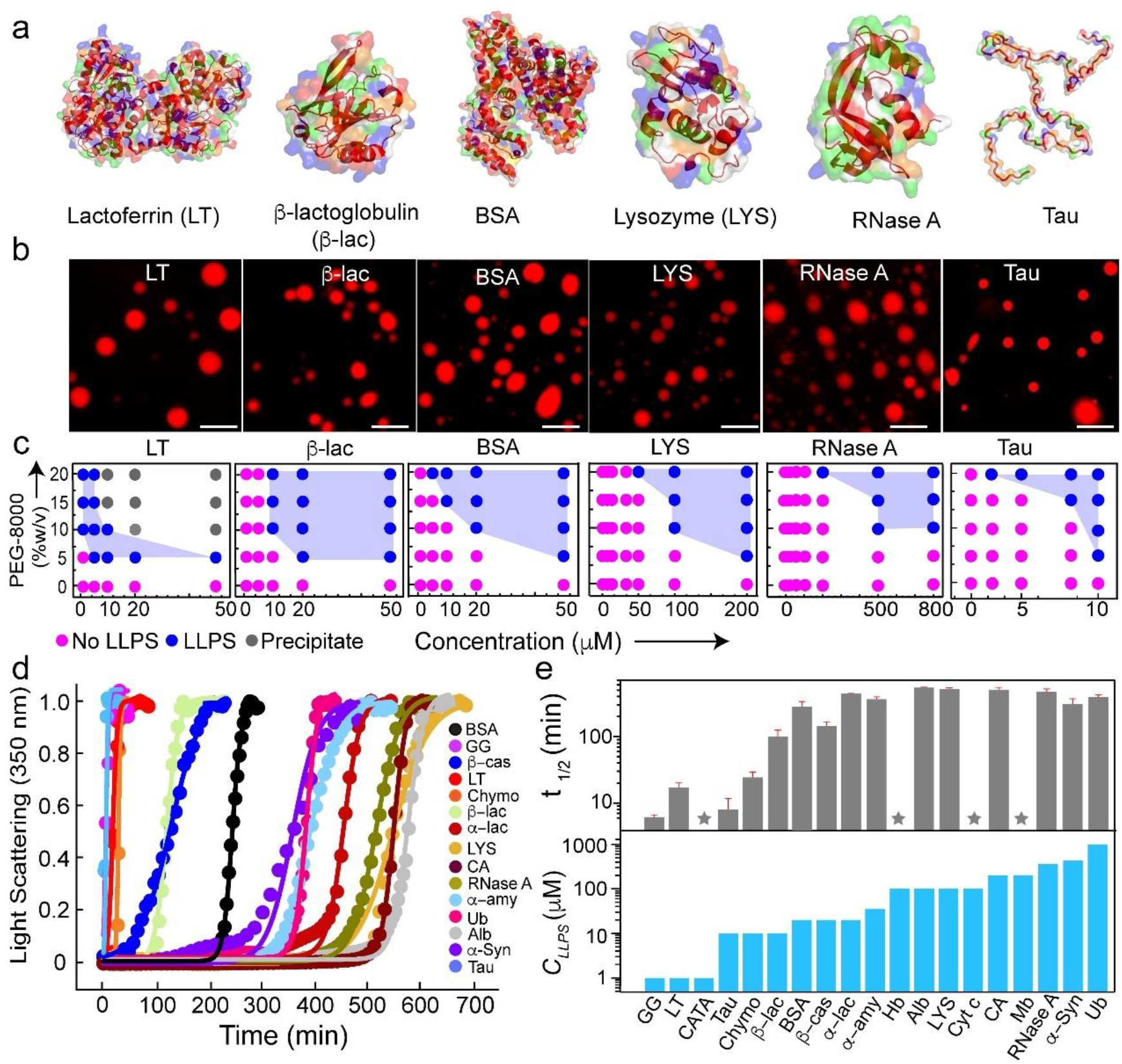
Liquid-liquid phase separation of various proteins *in vitro*. (a) Three-dimensional surface representation of selected proteins [LT (PDB ID: 1B0L), β-lac (PDB ID: 1QG5), BSA (PDB ID: 3V03), LYS (PDB ID: 1REX), RNase A (PDB ID: 1FS3), and Tau (PED00017e001)] with their embedded secondary structures (dark red). Positive, negative, and hydrophobic amino acids are represented in blue, red, and green colors, respectively. (b) Fluorescence microscopy images showing LLPS of selected NHS-Rhodamine labeled [10% (v/v) labeled to unlabeled] proteins (LT, β-lac, BSA, LYS, RNase A, and Tau) proteins in presence of 10% (w/v) PEG-8000. The samples were prepared in 20 mM sodium phosphate buffer (pH 7.4). Representative images are shown (*n=3*, independent experiments). The scale bar is 5 μm. (c) Phase regime of selected proteins (LT, β-lac, BSA, LYS, RNase A, and Tau) depicting LLPS at varying protein and PEG-8000 concentrations. The different states are represented with various color codes. The pink color indicates no LLPS (soluble state), the blue color indicates LLPS (condensate state), and the grey color indicates precipitation. The experiment was performed three independent times with similar observations. (d) Static light scattering experiment (at 350 nm) showing the kinetics of protein condensate formation with time at their *C*_*LLPS*_ and in presence of 10% (w/v) PEG-8000 (*n*=*2*, independent experiments). The light scattering values were normalized to 1. (e) ***Top panel***: t_1/2_ values depicting protein condensate kinetics determined from the sigmoidal fit from Fig. 1d. The data represent the mean ± s.d. for *n=2* independent experiments. ***Bottom panel***: *C*_*LLPS*_ of all proteins determined from the microscopic observation are represented with bar graphs. PEG-8000 was kept constant [10% (w/v)] to obtain a comparative measure of *C*_*LLPS*_ of all the proteins. The star symbol represents chromophore-containing proteins for which the light scattering experiment was not performed. The Y axis values are in log scale. *n=3* independent experiments.

IUPred2A^49^, SMART^50^, and CatGranule^51^, for predicting the presence of IDRs, LCDs, and their LLPS propensities, respectively. Our data revealed that a subset of proteins possess LLPS propensity as well as sequence/s featuring intrinsic disorders/LCD regions. On the other hand, many proteins such as lysozyme (LYS) and β-lactoglobulin (β-lac) did not exhibit any such feature (Fig. S2 and Table S2). To test whether these proteins can undergo LLPS *in vitro*, we purified all the proteins using size exclusion chromatography (SEC) and examined for LLPS using fluorescence microscopy (by labeling the proteins with NHS-Rhodamine) in the presence of PEG-8000 (Fig. 1b and S3). To construct the LLPS regimes, proteins at varying concentrations were incubated with different concentrations of PEG-8000 at physiological pH 7.4. We observed that all proteins could undergo LLPS at different concentrations, thereby exhibiting a varied phase space (Fig. 1c and S4a). The observed LLPS of proteins was not due to protein degradation as evident from the protein band/s in SDS-PAGE after LLPS (Fig. S5).

From this phase space, we further determined the critical concentration (estimated minimum concentration required for LLPS) (*C*_*LLPS*_) of the proteins in the presence of 10% (w/v) PEG-8000. Proteins such as lactoferrin (LT), γ-globulin (GG), and catalase (CATA) required as low as 1 μM concentration to undergo LLPS, while ubiquitin (Ub), and RNase A required a very high protein concentration (≥500 μM) for phase separation (Fig. S4a). To further evaluate the kinetics of LLPS, we performed a static light scattering experiment (at 350 nm) with their respective *C*_*LLPS*_ in the presence of 10% (w/v) PEG-8000 over time (Fig. 1d). Similar to their phase behavior, the data revealed that kinetics of LLPS also varied across different proteins (Fig. 1d and 1e). At the end of the light scattering experiments, the condensate formation was verified using differential interference contrast microscopy (DIC) (Fig. S4b). The light scattering data were fitted with a sigmoidal growth kinetics model (see method section) and the t_1/2_ for LLPS was determined for all the proteins (Fig. 1e). The data revealed that proteins with low *C*_*LLPS*_ exhibited faster LLPS kinetics in contrast to proteins with higher *C*_*LLPS*_. Overall, the data provide promising evidence in support of LLPS being a generic phenomenon of proteins.

### Liquid-like property of the phase separated condensates

Typical characteristics of phase separated condensates include condensate fusion upon contact, temperature reversibility, and rapid fluorescence recovery after photobleaching (FRAP). To examine the dynamic nature of the molecules inside the condensates, we performed FRAP using 10% (v/v) rhodamine-labeled proteins. At the initial time of condensate formation (0 h), most of the proteins showed rapid recovery of fluorescence (∼80-100% recovery) with a short half-life (t_1/2_) (<5 s); while a few proteins showed partial recovery (e.g., LT and GG showed 50-60% recovery) with higher t_1/2_ values (>10 s) (Fig. 2a-c and S6a). We hypothesized that extensive intermolecular interactions might result in their partial solid-like behavior leading to reduced fluorescence recovery (also supported by their very low *C*_*LLPS*_). The liquid-like property of the condensates was further supported by fusion events (Fig. 2d and S6b) and dissolution of condensates upon increased temperature (at 45 °C). The protein condensates however reappeared upon incubating back to 37 °C (Fig. 2e and S6c), suggesting their thermo-reversible property. To examine whether LLPS is associated with the conformational transition of the proteins, we characterized the protein structure using circular dichroism (CD) spectroscopy. We observed no substantial change in secondary structure/s upon phase separation (Fig. 2f and S7), which was also confirmed by FTIR analysis (Fig. S8 and S9). To analyze the morphology of liquid condensate, various protein LLPS samples were examined through transmission electron microscopy (TEM), which mostly showed circular protein-rich condensates (Fig. 2g and S6d). The data therefore suggest that proteins can form thermo-reversible, liquid condensates without significant alteration in their secondary structures.

**Figure 2.**
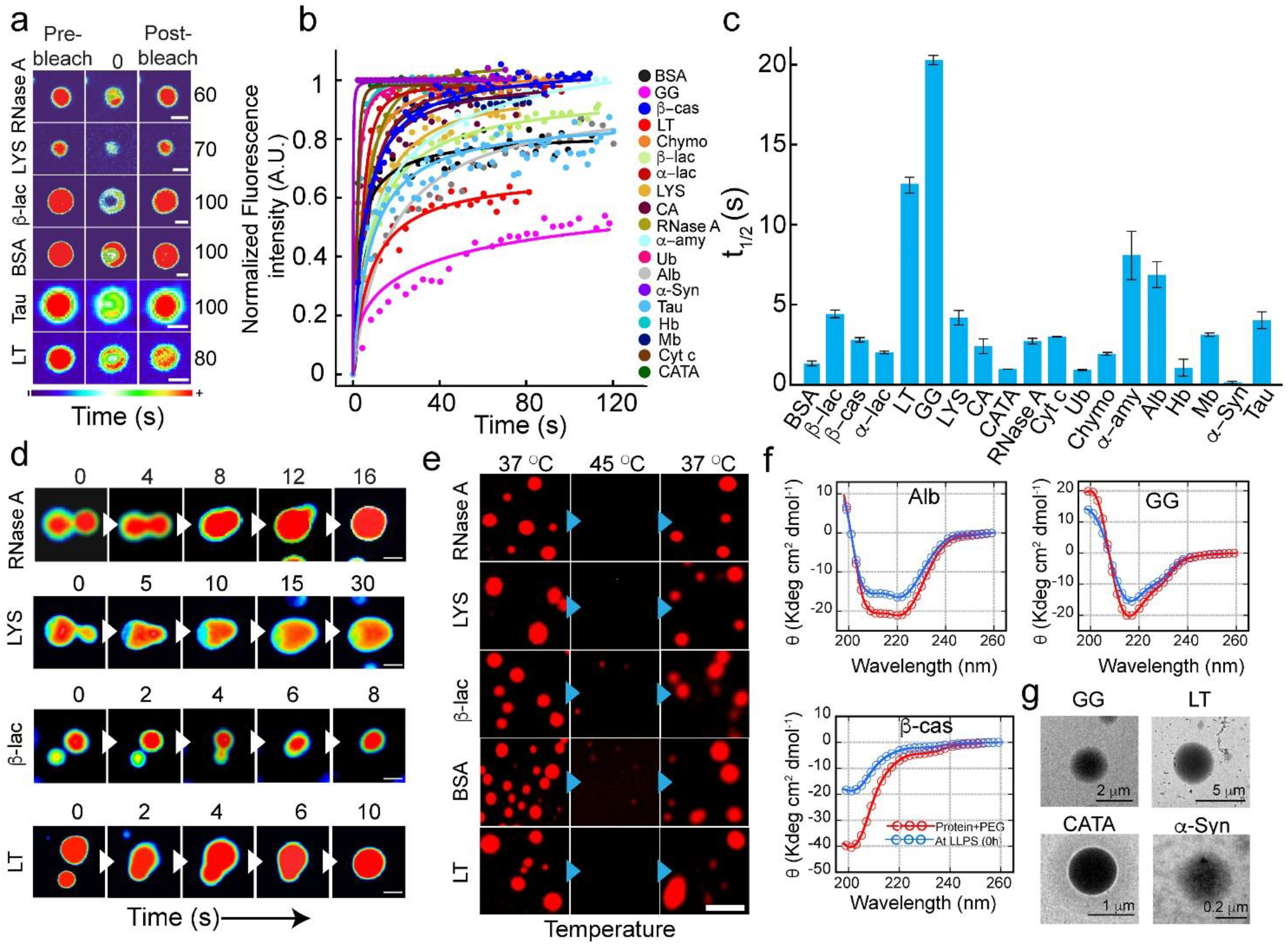
Liquid-like properties of the various protein condensates. (a) Representative images showing the liquid condensates (immediately after formation, 0 h) during FRAP (before bleaching, at bleaching, and after bleaching) for selected proteins (RNase A, LYS, β-lac, BSA, Tau, and LT). The scale bar is 2 μm. The images are represented in the ‘thermal’ lookup table (LUT) for better visualization. (b) Normalized FRAP curves obtained for all the protein condensates at 0 h (immediately after LLPS) are plotted against time. *n=3* independent experiments were performed. (c) The bar plot of t_1/2_ values showing fluorescence recovery after photobleaching of protein condensates at 0 h. The data represents the mean ± s.e.m. for *n=3* independent experiments. (d) Time-lapse images showing fusion events of condensates formed by selected proteins over time (RNase A, LYS, β-lac, and LT). Images are represented in ‘royal’ LUT for better visualization. Representative results are shown. The scale bar is 5 μm. The experiment was repeated two times. (e) Fluorescence microscopy images showing thermo-reversibility (37 °C → 45 °C → 37 °C) of selected NHS-Rhodamine labeled [10% (v/v)] protein condensates formed at their respective *C*_*LLPS*_ in the presence of 10% (w/v) PEG-8000 (at 0 h). Representative images are shown. The experiment was performed two times with similar observations. The scale bar is 5 μm. (f) CD spectroscopic analysis of selected proteins (Alb, β-cas, and GG) showing no substantial changes in the secondary structural conformation of the proteins upon phase separation. The red and blue color indicating protein CD spectra before and after phase separation in the presence of PEG-8000 [10% (w/v)]. (g) Representative TEM images showing the morphology of protein condensates formed by GG, LT, CATA, and α-Syn. *n=2* independent experiments were performed.

### Liquid-to-solid transition of protein LLPS

Liquid-to-solid phase transition of protein condensates is often associated with toxic amyloid fibril formation in various neurodegenerative diseases such as ALS, AD, and PD^17-20^. However, such viscoelastic transition can also help in various cellular functions^5,6,52^ including oocyte dormancy (Balbiani body^53^) and heterochromatin assembly^54-56^. We wanted to investigate whether the condensates formed by the various proteins in our study also undergo solidification with time. We incubated various protein condensates for 48 h (at 37 °C) and performed FRAP and temperature reversibility (Fig. 3a-d and S10a, b) studies. FRAP analysis of condensates at 48 h revealed substantially slower recovery (higher t_1/2_) for most of the proteins compared to freshly formed liquid condensate (0 h) (Fig. 3c). Intriguingly, a few proteins (GG, LT, Tau, α-Syn, β-cas, and CATA) did not recover after photo-bleaching at 48 h (Fig. 3c), suggesting their transition to a solid-like state. This was also consistent with the thermo-reversibility study as these condensates did not dissolve upon increasing the temperature to 45 °C (Fig. 3d and S10b). To examine the possible structural changes due to liquid-to-solid transition, we performed CD spectroscopy for a subset of proteins (which showed negligible fluorescence recovery after 48 h). We observed a significant decrease in molar ellipticity (θ) values after the liquid-to-solid transition by these proteins (Fig. 3e and S11). To further examine whether the decrease in molar ellipticity of proteins is associated with any structural changes or due to decrease in protein solubility (liquid to solid transition), we performed solid-state FTIR spectroscopy (Fig. S8 and S9). The data suggest that except for α-Syn, rest of the proteins did not undergo substantial structural transition during liquid to solid transition, consistent with CD data. Moreover, a subtle structural alteration was also observed for LT and β-cas and α-Syn showed major structural change from RC to β-sheet transition in FTIR spectroscopy (Fig. S8 and S9). To examine whether the liquid-to-solid transition by any of the proteins were associated with amyloid fibril formation, we performed ThT (binds to amyloid aggregates) fluorescence assay^18^. That data suggests that except α-Syn (bind strongly with ThT as expected^18^), no other proteins undergoing liquid to solid transition showed any significant ThT binding (Fig. 3f). This indicates that either crystal-like native packing/protein vitrification and/or amorphous aggregation might result in their solidification^1,2,6,22,57^. To further characterize the morphology of the condensates after 48 h, we analyzed the condensates using TEM (Fig. 3g and S10c). The TEM images of GG and LT condensates showed a multiphasic nature as evident from different electron-dense/sparse regions, indicating protein assembly in the condensate (Fig. 3g). We also observed aggregate-like morphology around α-Syn condensates as previously reported^18,39,58,59^ (Fig. 3g). The data suggest that partial or full solidification might occur for protein condensates upon ageing with or without structural transition. Further, the generic nature of solidification (partial/full) by many proteins indicate the life time and material property of protein condensates *in vivo* might also be tightly regulated for the native cellular functions. However, when demixing ability of a cell is compromised, it would lead to toxic protein aggregation, leading to diseases_1,4,18-20_.

**Figure 3.**
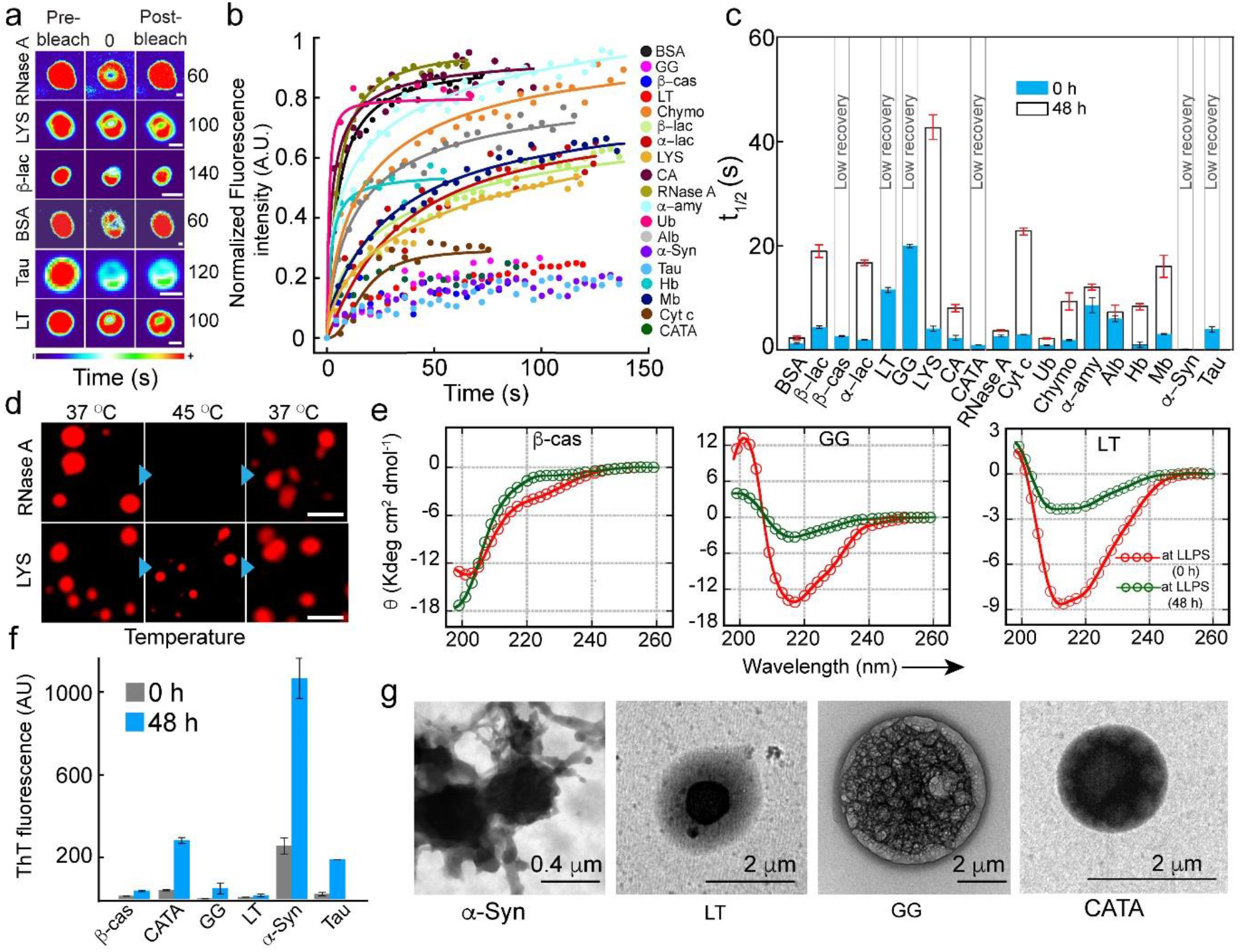
The liquid-to-solid transition of protein condensates. (a) Representative images showing selected protein condensates after LLPS (48 h) during FRAP analysis (before bleaching, at bleaching, and after bleaching). The images are represented in ‘thermal’ LUT for better visualization. RNase A and BSA show complete fluorescence recovery, whereas, LYS, β-lac, Tau, and LT show partial recovery. The scale bar is 2 μm. (b) Normalized FRAP profile of the phase separated condensates at 48 h by various proteins showing different fluorescence recovery. A subset of protein condensates after 48 h show reduced fluorescence recovery, indicating that they might undergo solidification over time. *n=3* independent experiments were performed. (c) The t_1/2_ values calculated from FRAP of all proteins at 0 h (blue) and 48 h (white), showing slow fluorescence recovery of protein condensates after (48 h). Notably, t_1/2_ values could not be calculated for β-cas, LT, GG, CATA, α-Syn, and Tau due to the negligible recovery after photobleaching. The data represent the mean ± s.e.m. for *n=3* independent experiments. (d) Fluorescence microscopy image of NHS-Rhodamine labeled condensates (10% v/v) by RNase A (48 h) showing thermo-reversibility upon heating (45 °C) and cooling (37 °C). The LYS condensates did not dissolve upon heating, suggesting liquid-to-solid phase transition after 48 h. The scale bar is 5 μm. Representative images are shown and the experiment was performed two times with similar observations. (e) CD spectra of selected proteins (LT, GG, and β-cas) immediately after LLPS (0 h, red) and after 48 h (green) demonstrating the secondary structure of the protein condensates over time (*n=2*, independent experiments). (f) ThT fluorescence intensity of different proteins immediately after LLPS (0 h) and after 48 h are shown. An increase in ThT intensity for α-Syn at 48 h suggests the formation of ThT positive, amyloid aggregates. The data represent the mean ± s.d. for *n=2* independent experiments. (g) TEM micrographs showing the appearance of multiphasic architectures by various protein condensate (LT, GG, and CATA), while amyloid fibril formation by α-Syn condensate. *n=2* independent experiments were performed.

### The charge distribution and exposed hydrophobic surface of a protein drive its LLPS

We hypothesized that proteins undergo LLPS through different intermolecular interactions based on their charge and hydrophobicity. This is due to the different structural fold/s and amino acid sequences of the proteins. According to the Flory Huggins theory^60^, the important criteria driving phase separation are (a) length of residues capable of intermolecular interactions (which is directly proportional to molecular weight (MW)) and (b) their respective interaction strengths. To establish a connection between protein sequence and its propensity of phase separation, we plotted various sequence-specific parameters with *C*_*LLPS*_ (Fig. 4 and S12). Although, achieving a perfect correlation between the sequence-specific quantities and *C*_*LLPS*_ is unlikely in this case as *C*_*LLPS*_ was estimated manually using the microscopic technique with an interval of concentrations (Fig. 1, S3 and S4). Further, LLPS is also likely to be governed by one or a combination of multiple intermolecular interactions (electrostatic, hydrophobic, and H-bonding) in proteins^18,40,61-63^. Indeed, we observed an approximate negative correlation (in log-log comparison) between *C*_*LLPS*_ and the absolute charge [√(N^2^ + P^2^)] of the protein (Fig 4a). Interestingly, the plot emerged with two distinct clusters, where one cluster (upper region) comprised high molecular weight (MW) proteins (MW > 40000 Da) and the other one (lower region) comprised proteins with low molecular weight (MW < 40000 Da). When we analyzed the two clusters separately, the proteins in each cluster however showed a weak correlation. This suggests that not only the absolute charge but also other factors such as hydrophobic interactions, H-bonding, and patterning/segregation of oppositely charged residues might also play a role in determining the *C*_*LLPS*_. To examine the role of hydrophobic interactions in proteins undergoing LLPS, we probed exposed hydrophobic surface with ANS binding study^64^. The data showed a significant increase in ANS fluorescence for BSA and a moderate increase for Alb, CATA, and β−lac, suggesting hydrophobic interactions might play a role for LLPS in these proteins. (Fig. 4b). We omitted these proteins while further computing the correlation to probe the role of electrostatic interactions. We also omitted α-Syn, as this protein is known to form LLPS through the hydrophobic NAC region at high concentration and longer incubation time or in presence of low pH where the hydrophobic region is exposed^18,39^. Once the proteins with significant ANS binding were omitted for analysis, a relatively strong correlation was emerged between absolute charge and *C*_*LLPS*_ for both the clusters (Fig. 4a) (R^2^=0.83 for higher MW cluster and R^2^=0.7 for lower MW cluster). We further computed a previously studied parameter kappa (κ)^65^ that accounts for the segregation of charges along the protein-polymer contour (Fig. S12a). As charge segregation might play a role for LLPS of proteins^66^ and the phase separation behavior would depend both on the strength of the interactions and the number of amino acids (length of the sequence), we plotted κ_*_C, which is the product of kappa (κ) and the total number of charged amino acids (C). In contrast to κ, κ_*_C showed a correlation with *C*_*LLPS*_ (Fig. 4c) similar to absolute charge (Fig. 4a), resulting in two clusters of proteins based on MW. Strikingly in this case, the slope of negative correlation was less steep for the low MW protein cluster. This suggests that for low MW proteins, change in charge segregation alter the propensity of LLPS to a greater extent, in comparison to the high MW proteins. In the above correlation studies, Mb was omitted since it appeared as an outlier in the both the cases. Similar to charge, *C*_*LLPS*_ also showed a negative linear correlation (in semi-log comparison) with the number of aromatic residues, (Fig. S12b), and the number of positive plus aromatic residues (Fig. S12c), suggesting LLPS is also driven by cation-π interactions within the protein^23,26,30^.

**Figure 4.**
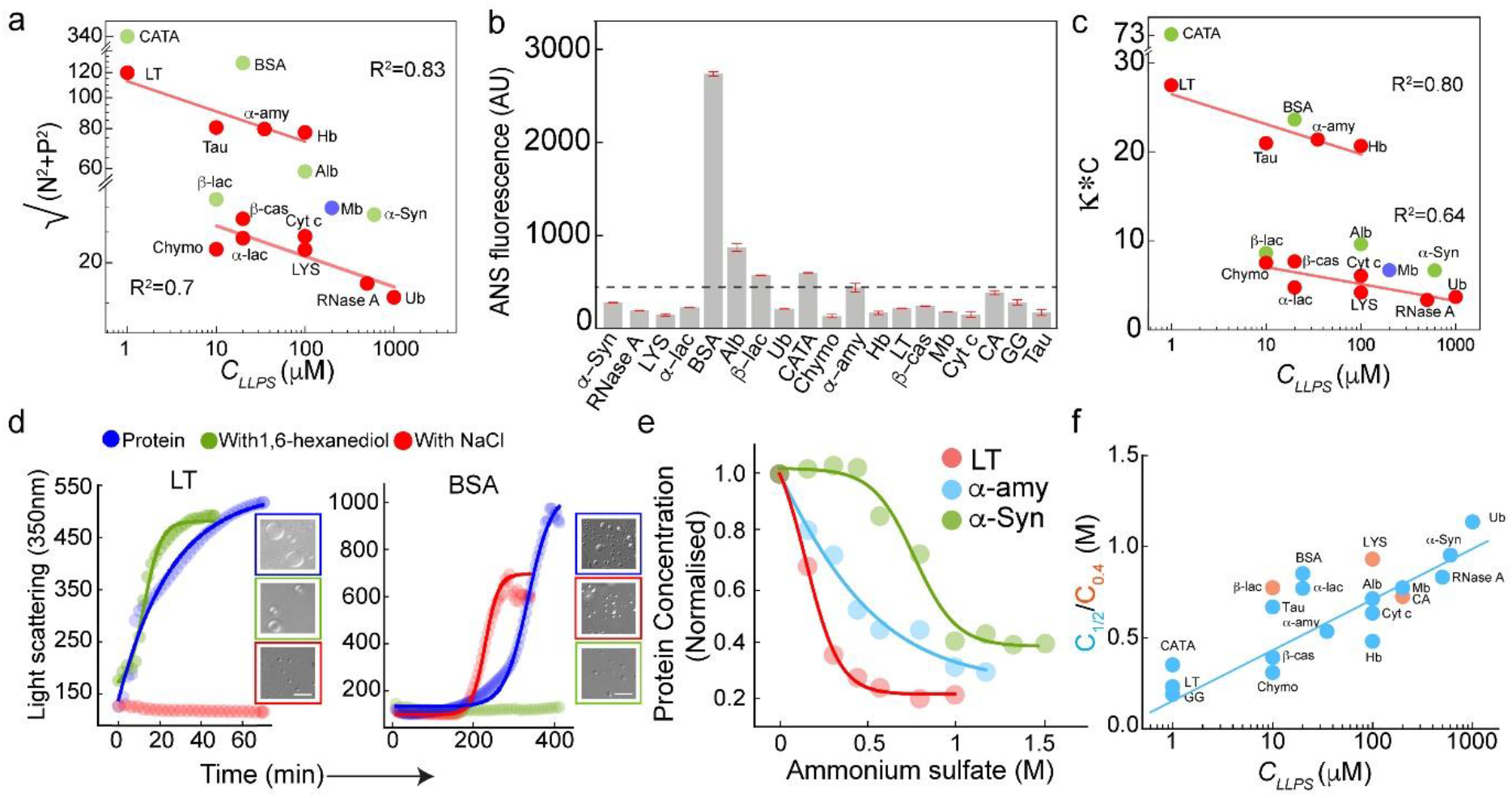
Intermolecular interactions govern LLPS of all proteins. (a) Correlation plot of absolute charge [√(N^2^+ P)^2^] and *C*_*LLPS*_ (in log-log scale) showing segregation of proteins into two clusters based on the molecular weight (MW) [upper region (MW > 40000 Da) and lower region (MW < 40000 Da)]. Straight lines represent correlation (R^2^ = 0.83 and 0.7 for high and low MW clusters, respectively) in the absence of respective outliers. Red color represents the proteins in which LLPS is largely driven by electrostatic interactions, green color represents the outliers (hydrophobic interactions) and blue color represents Mb, which is omitted from the fitting analysis. (b) ANS fluorescence of all proteins showing the extent of exposed hydrophobic surface. The data represent the mean ± s.d. for *n=2*, independent experiments. (c) κ*C, the product of Kappa and the number of charged amino acids showing a negative linear correlation (in semi-log scale) with the *C*_*LLPS*_. Red color represents the proteins in which LLPS is largely driven by electrostatic interactions, green color represents the outliers (hydrophobic interactions) and blue color represents Mb, which omitted from the fitting analysis. (d) The light scattering measurement at 350 nm showing relative inhibition of LLPS (LT and BSA) in the presence of either 150 mM NaCl (red) or 10% (w/v) 1, 6-hexanediol (green). All the LLPS experiments were performed in the presence of 10% (w/v) PEG-8000. Blue color indicates light scattering measurement of protein in presence of 10% (w/v) PEG-8000 only. Representative DIC images showing absence or presence of condensate formation in light scattering experiments. The scale bar is 5 μm, *n=2*, independent experiments were performed. (e) The change in soluble protein concentrations after each addition of ammonium sulphate showing the decrease in protein concentration with an increased concentration of ammonium sulfate (*n=2*, independent experiments). *C*_*1/2*_ was determined from the concentration decay curve with curve fitting. (f) Correlation plot of *C*_*1/2*_ and *C*_*LLPS*_ (in semi-log scale) representing proteins with low *C*_*LLPS*_ exhibited precipitation at a minimum ionic strength of ammonium sulfate and vice versa. The straight line in blue is just a guide to an eye (*n=2*, independent experiments). For LYS, CA and β-lac (orange), *C*_*0*.*4*_ was calculated instead of *C*_*1/2*_ because in the present experimental condition, 50% protein precipitation could not be reached.

After predicting that proteins might mostly use either electrostatic or hydrophobic (or in combination) interactions for LLPS, we examined the kinetics of LLPS for selected proteins in the presence of salt, NaCl (to disrupt electrostatic interaction)^13,31^ or 1,6-hexanediol (to disrupt hydrophobic interaction)^61,67^ using static light scattering (at 350 nm) (Fig. 4d and S13a). Our data showed that LLPS of LT and Chymo (as predicted electrostatic interaction for LLPS) was largely inhibited by the addition of 150 mM NaCl; while there was no effect due to the presence of 10% (w/v) 1,6-hexanediol. In contrast, LLPS of BSA and Alb (with ANS binding due to exposed hydrophobic surface) was substantially inhibited by presence of 10% (w/v) 1,6-hexanediol, but no difference in scattering intensity was observed in presence of 150 mM NaCl (Fig. 4d and S13a).

To further examine the role of protein solubility in determining the *C*_*LLPS*_, proteins were subjected to the increasing concentrations of ammonium sulfate for precipitation^68^ and the remaining soluble protein concentration was plotted against ammonium sulfate concentration (Fig. 4e and S13b). In this context, it has been previously shown that LLPS of FUS protein was greatly inhibited by increasing its solubility with solubility tag^69^. Further, proteins with lesser solubility are known to possess higher propensity for self-assembly, protein aggregation and LLPS^11,35,68,70^. Indeed, when *C*_*1/2*_ (concentration of ammonium sulfate required for 50% protein precipitation) was plotted against *C*_*LLPS*_, we observed an approximate positive correlation suggesting solubility of protein dictates their *C*_*LLPS*_ (Fig. 4f). The data suggest that solubility and LLPS of proteins might be majorly governed by the extent of their intermolecular interactions.

### LLPS by charged and neutral polypeptides

We hypothesized that if high intermolecular interactions (*via* hydrophobic/electrostatic) are the only necessary prerequisites for phase separation, then even small polypeptides at optimum concentration can undergo LLPS. To examine this, we designed four 10-residue polypeptides [(Gly)_10_, (Val)_10_, (Arg)_10_, and (Asp)_10_] and characterized them using MALDI and LC-MS (Fig. 5a and S14). We speculated that the non-polar polypeptide, (Gly)_10_, would require a very high concentration for LLPS due to lack of polyvalency/side chains by simplest amino acid, glycine^26^. However, (Val)_10_ might undergo intermolecular interaction based on hydrophobic interactions, which will facilitate its LLPS. Interestingly, both the peptides showed LLPS at high concentrations in 20 mM sodium phosphate buffer, pH 7.4 in the presence of 10% PEG. The (Gly)_10_ showed LLPS ≥ 2 mM concentration, while (Val)_10_ showed LLPS ≥ 1 mM concentration. This suggests (Val)_10_ possess higher LLPS propensity compared to (Gly)_10_ (Fig. 5b, c, and S15a) due to its hydrophobic side chains. In contrast to (Gly)_10_/(Val)_10_, the liquid condensate formation of charged homo-polymers might occur upon neutralization of charged residues^25,33^. Indeed, our data showed that (Arg)_10_ and (Asp)_10_ formed condensates in the presence of 4 M NaCl [with 10% (w/v) PEG-8000] at a concentration of ≥ 2 mM and ≥ 8 mM, respectively (Fig. 5b, c, and S15a-b). This data suggests that at charged neutralized state, poly arginine might possess higher polyvalency for LLPS in comparison to poly aspartic acid. As expected, in absence of NaCl, we did not observe any phase separation by these polypeptides due to their charge-charge repulsion. Here, the presence of 10% (v/v) labeled polypeptide did not affect the condensate formation by both of the charged polypeptides (Fig. S15c).

**Figure 5.**
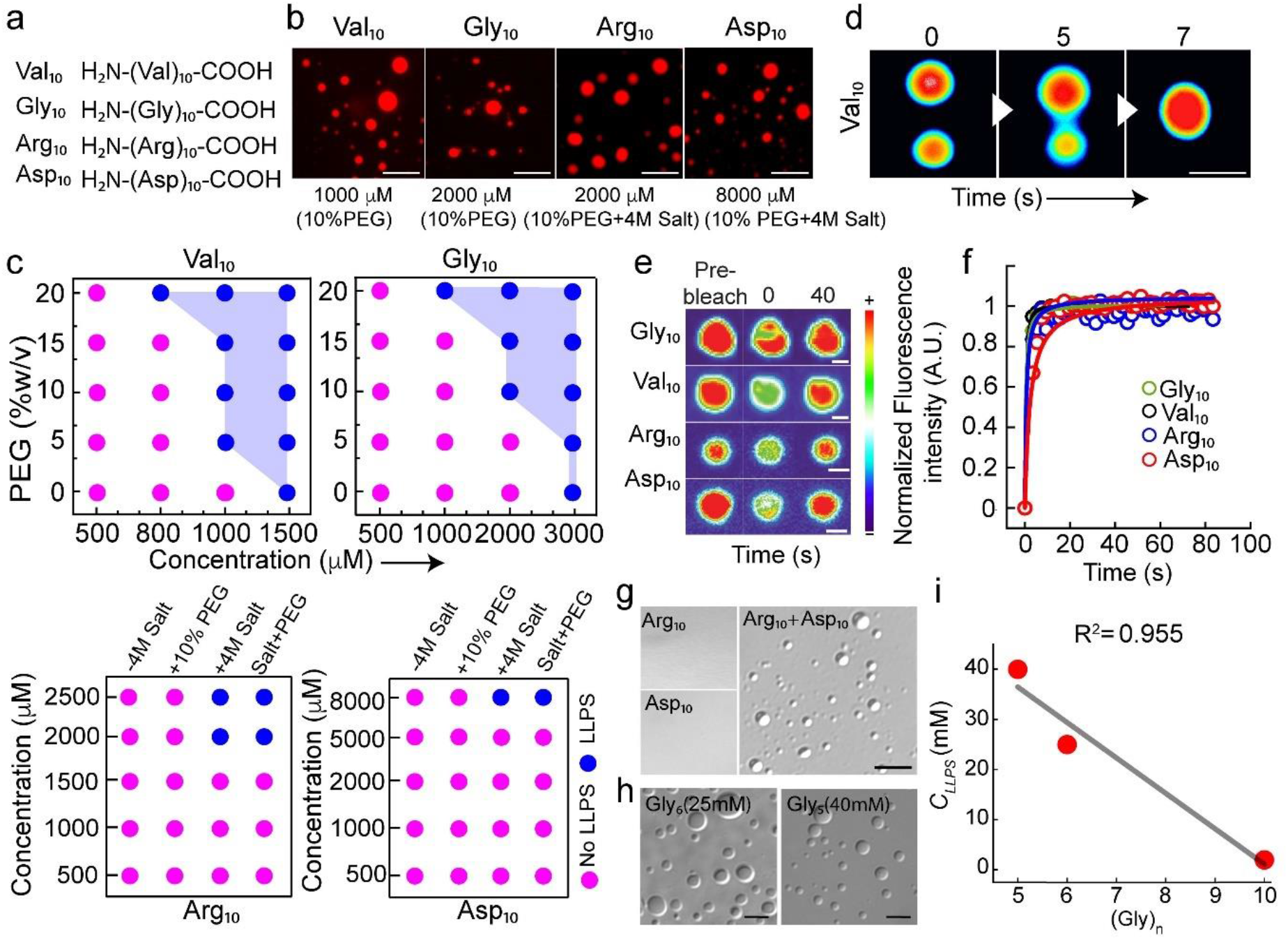
Liquid-liquid phase separation of polypeptides. (a) Sequence information of designed polypeptides with ten residues are depicted. (b) Fluorescence microscopy images showing liquid condensate formation by NHS-Rhodamine-labeled (10% v/v) polypeptides at their respective *C*_*LLPS*_ in the presence of 10% (w/v) PEG-8000. Notably, (Arg)_10_ and (Asp)_10_ formed liquid condensates in presence of 4 M NaCl (*n=2*, independent experiments). The scale bar is 5 μm. (c) ***Top Panel***: LLPS regimes of (Val)_10_ and (Gly)_10_ at varying polypeptide and PEG-8000 concentrations are shown. ***Bottom Panel:*** LLPS regime of (Arg)_10_ and (Asp)_10_ in the presence and absence of 4 M NaCl and 10% (w/v) PEG-8000 at varying peptide concentrations are shown. The pink color indicates no LLPS (soluble state) and the blue color indicates LLPS (condensate state). (d) Time-lapse images of (Val)_10_ condensate showing fusion event (represented in ‘royal’ LUT for better visualization). The scale bar is 5 μm (*n=2*, independent experiments). (e) Representative FRAP images (before bleaching, at bleaching, and after bleaching) of polypeptide condensates at 0 h (represented in ‘thermal’ LUT). The scale bar is 2 μm. (f) Normalized FRAP profile showing complete fluorescence recovery of the phase separated condensates (at 0 h) of polypeptides (Gly)_10_, (Val)_10_, (Arg)_10_, (Asp)_10_, represented in green, black, blue, and red color, respectively (*n=3*, independent experiments). (g) DIC microscopy images of (Arg)_10_ and (Asp)_10_ condensates at their respective *C*_*LLPS*_ when mixed (in absence of salt) are shown. Respective polypeptides at *C*_*LLPS*_ without mixing (in absence of salt) were used as controls. The scale bar is 5 μm. The experiment was repeated three times with similar observations. (h) DIC microscopy images of (Gly)_5_ and (Gly)_6_ showing condensate formation at their respective *C*_*LLPS*_ in presence of 10% (w/v) PEG-8000 (*n=3*, independent experiments). (i) A correlation plot (with R^2^ value: 0.955) of *C*_*LLPS*_ and length of (Gly)_n_ polypeptides suggesting that as the polypeptide length increases, the *C*_*LLPS*_ decreases linearly. The experiment was repeated twice with similar observations.

The LLPS by these homo-polypeptides was also further characterized by fusion (Fig. 5d and S15d) as well as FRAP studies (Fig. 5e, f and S15e-f). Both (Gly)_10_ and (Val)_10_ condensates showed fusion upon contact to form larger condensates and all the polypeptide condensates showed complete fluorescence recovery after photobleaching, confirming their liquid-like property. Further, morphology of the condensates was examined using TEM, which revealed homogeneous electron density of the condensate state of these polypeptides (Fig. S15g). The data therefore suggest that small homo-polypeptide also undergo LLPS but with relatively high *C*_*LLPS*_ suggesting that intermolecular interaction between these polypeptides is much less prevalent compared to large proteins. To further examine whether intermolecular interactions between oppositely charged polypeptide facilitate LLPS, we examined the co-LLPS of (Arg)_10_ and (Asp)_10._ When two oppositely charged polypeptides were mixed, we observed spontaneous phase separation in absence of salt, suggesting charge neutralization favors their co-LLPS (Fig. 5g and S15h). In identical condition, however, the individual polypeptide did not show any LLPS (Fig. 5g and S15h). To further examine the contribution of polypeptide length in *C*_*LLPS*_, we chose glycine polypeptides. When LLPS study was performed with (Gly)_6_ and (Gly)_5_ in 20 mM sodium phosphate buffer, pH 7.4 in the presence of 10% PEG, we observed that (Gly)_5_ and (Gly)_6_ required 40 mM and 25 mM concentration for their LLPS (Fig. 5h and S15i). Overall, the polymer length and *C*_*LLPS*_ of glycine polypeptides showed a negative linear correlation, suggesting that decrease in polypeptide length will increase the *C*_*LLPS*_ and vice versa (Fig. 5i).

### Heterotypic co-LLPS of various proteins in combination

Heterotypic phase separation by multiple proteins is implicated in many biological processes associated with membraneless organelle formation in cells^6,37,40,71^. In certain heterotypic/multicomponent phase separation system, a ‘scaffold’ protein is proposed to maintains the structural and thermodynamic integrity of the LLPS state, which further recruits various ‘client’ proteins for functionality^72,73^. Although, in this heterotypic co-LLPS process, whether any sequence preference exist between interacting proteins is not clear yet. To examine heterotypic phase separation by various combination of protein pairs, we investigated co-LLPS of BSA (FITC labeled) with other different proteins (NHS-Rhodamine labeled) in 20 mM sodium phosphate buffer, pH 7.4 in presence of 10% PEG. For experimental feasibility and to avoid large sampling (^19^C_2_, i.e., 171 possible combinations), we chose BSA as one of the representative partners and each of the other proteins as the second one. We observed that BSA undergo co-LLPS with each of the other proteins under study. Further, when both the proteins were mixed at their respective *C*_*LLPS*_ or at equal protein concentrations, they resulted in the formation of colocalized condensates (co-LLPS) (Fig. 6a, S16, S17, and S18). Interestingly, in some cases, heterotypic LLPS is favored even when half the respective *C*_*LLPS*_ of the proteins was mixed (Fig. 6f, S16, and S18), but no condensate formation was seen for the individual proteins without mixing. The data suggest that the presence of other proteins not only increased the overall crowding but also favored the heterotypic interaction for co-LLPS where BSA acts as one of the partners. This may be due to the fact that BSA possess exposed hydrophobic surface as well as high absolute charge for intermolecular interactions (Fig. 4) facilitating co-LLPS with other proteins. To examined further the co-LLPS with different partners, we randomly chose other protein pairs (Table S3) for study using their respective *C*_*LLPS*_. Although in some combination of protein pairs, co-LLPS (yellow colocalization) was observed, while for others, either one protein phase separated or LLPS was completely inhibited by both the partners in mixture (Fig. S19 and S20). The data suggest that the heterotypic co-LLPS by the different proteins is majorly dictated by their effective protein-protein interactions

**Figure 6.**
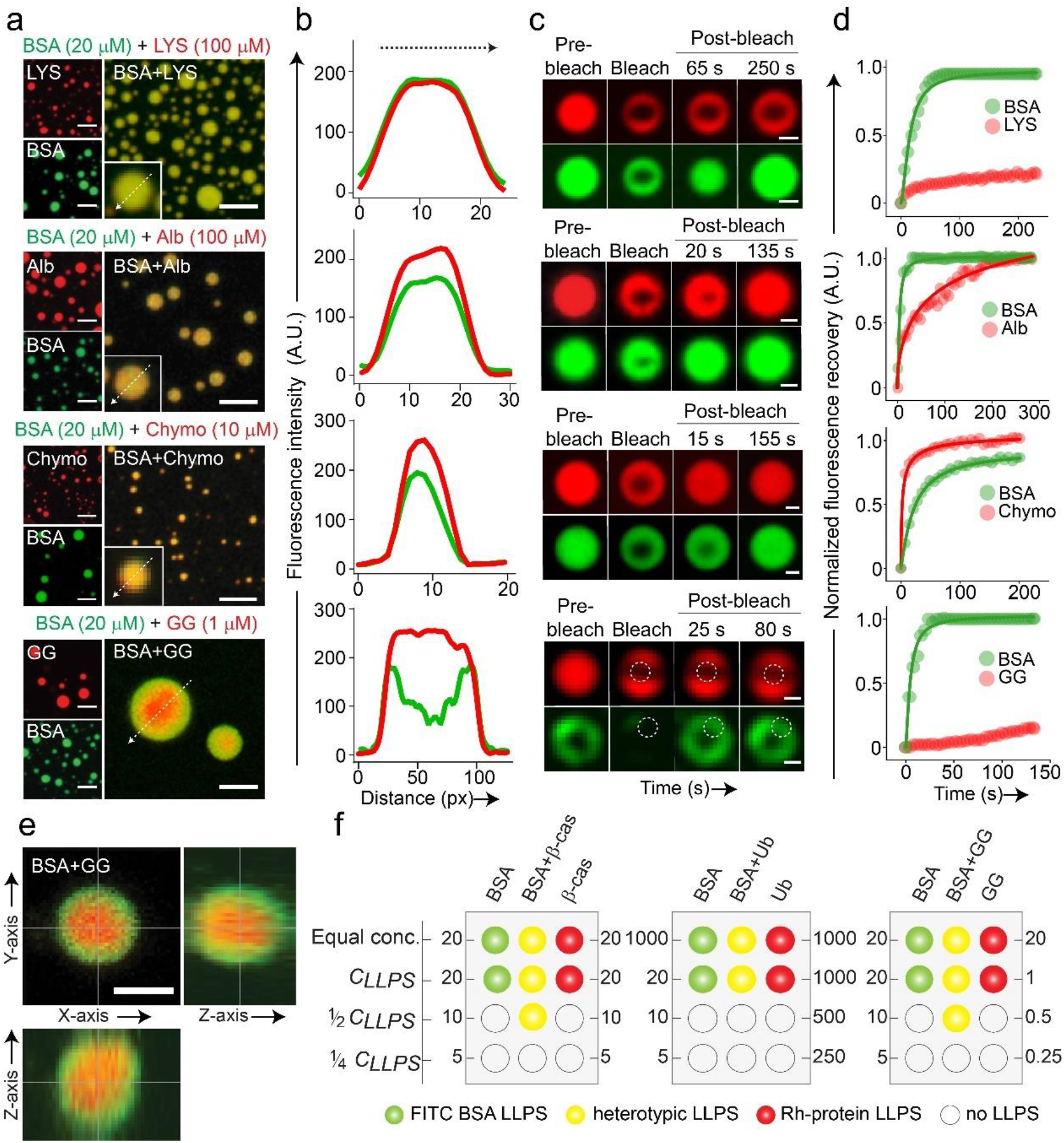
Heterotypic liquid condensate formation by various combinations of proteins. (a) Representative fluorescence microscopic images showing heterotypic LLPS of FITC labeled BSA and NHS-Rhodamine labeled LYS, Alb, Chymo, and GG when mixed at their respective *C*_*LLPS*_. Respective labeled proteins alone (at *C*_*LLPS*_) were used as control, which is shown on the left-hand side of each image. The scale bar is 5 μm. The experiment was repeated twice with similar observations. (b) Fluorescence intensity profile of heterotypic phase separated condensates showing co-localization of FITC labeled BSA and NHS-Rhodamine labeled LYS, Alb, Chymo, and GG inside the same condensate. A red core-like structure was seen in BSA/GG system due to the preferential segregation of GG. (c) Representative FRAP images at respective fluorescent channels (before bleaching, at bleaching, and after bleaching) of the heterotypic condensate showing different recovery kinetics depending on the proteins mixed in the system. The FITC labeled BSA is represented in green color and the NHS-Rhodamine labeled partner proteins are represented in red color. The scale bar is 2 μm (*n=2*, independent experiments). (d) Normalized FRAP profile of the heterotypic condensates showing different fluorescence recovery of the individual protein component in the two-protein LLPS system. (e) The representative orthogonal Z-sectioning of the FITC labeled BSA and NHS-Rhodamine labeled GG heterotypic condensate confirming the preferential segregation of GG at the center of the heterotypic condensate. The panel on the right side showing the YZ image axis and the bottom panel showing the XZ image axis of the condensate. The scale bar is 2 μm. The experiment was repeated twice with similar observations. (f) Schematic representation of two-protein component system and corresponding single-protein component showing concentration-dependent heterotypic liquid condensate formation. The green color indicates LLPS of FITC labeled BSA only (single-component), red color indicates LLPS of NHS-Rhodamine labeled protein (single-component) and yellow color indicates heterotypic co-LLPS. The empty symbol indicates no LLPS event.

Furthermore, in the case of GG/LT co-LLPS with BSA, we observed the appearance of GG or LT-rich, core-like multiphasic structure (Fig 6a-b, e and S16), which suggests the high preferential segregation of GG and LT resulting their sub-compartmentalization in heterotypic co-LLPS^1,6,40^. Similar observation was also obtained for RNase A/GG pair, where GG showed core-like segregation at the center of heterotypic condensate (Fig. S20). Since GG and LT showed rapid liquid-to-solid transition (Fig. 2b), the data therefore suggest that faster solidification of one partner might be the cause for its segregation inside the condensate. This is also consistent with FRAP data of GG/BSA heterotypic condensate, where GG showed negligible fluorescence recovery upon photobleaching compared to BSA, immediately after condensate formation (Fig. 6c, d).

Further, FRAP data showed for BSA/LYS, BSA/Chymo and BSA/Alb heterotypic condensates, the fluorescence recovery of LYS, BSA and Alb showed slower recovery (higher t_1/2_), respectively compared to their counterpart in the heterotypic condensate (Fig. 6c-d and S21). This data suggests that even though the condensate formation happens due to intermolecular interactions in the heterotypic systems, the dynamic nature of individual protein component is substantially different in the heterotypic condensates.

Taken together, our data suggest that depending upon the protein pair, and their respective CLLPS, the heterotypic condensates formation and their further solidification might be modulated. Moreover, the relative rate of liquid-to-solid transition of each component can modulate their spatial distribution inside the multiphasic condensates.

## Discussion

LLPS of biomolecules such as proteins and RNA has emerged as an important phenomenon responsible for many biological processes in cells^1-6^. Previous studies have shown that IDRs, LCDs, PLDs, and peptides/proteins with spacer and sticker motifs are critical determinants for protein LLPS^23-27,29^. Although these criteria might be facilitating the multivalent interactions necessary for protein assembly on a physiologically relevant scale; they might not be mandatory for LLPS in general. We hypothesized that LLPS might be a common property of proteins and polypeptides under specific conditions (such as increased concentration and crowding) where the threshold extent of intermolecular interactions can cross the energy barrier for LLPS^11,32,36,37,44^. When a polypeptide folds into a native 3-dimensional structure in an aqueous solution, the hydrophilic amino acids generally constitute the surface of the protein and the hydrophobic amino acids reside within the folded core of the protein. Therefore, the protein surface-mediated interactions with water are the major driving force for the protein to be soluble. Apart from solubility, electrostatic and H-bonding also can be attributed to intermolecular interactions leading to LLPS when proteins are in close proximity. In addition to electrostatic interactions, exposed hydrophobic surface (which otherwise is inaccessible in folded protein) is also highly susceptible for intermolecular interactions and could drive protein LLPS^18,61-63^. Indeed, we observed that various proteins with diverse sequences and structures can readily undergo LLPS, however with a wide variation in their *C*_*LLPS*_ and t_1/2_ (Fig. 1). *C*_*LLPS*_ and t_1/2_ of observable liquid condensate formation not only depends on the intrinsic protein sequence/structure but also extrinsic factors such as pH, presence of salts, and other cellular factors^10,13,31,38^. For example, α-Syn undergoes LLPS at a very high concentration (600 μM) in salt-free buffer systems. However, increasing the salt concentration or lowering the pH drastically brings down the *C*_*LLPS*_ of α-Syn LLPS due to charge neutralization at the C-terminus^18,39^.

Interestingly, the phase separation by all these proteins does not require a misfolding or drastic structural transition, suggesting that low solubility or high enough concentration is sufficient for inducing LLPS (Fig. 2 and 4). Indeed, many functional proteins in the human proteome are very close to their solubility limit and slight fluctuations in protein concentration/cellular environment might promote their transition from soluble phase to a condensate state^32,35,44,45^. Consistent with all previous studies, most LLPS systems maintain their liquid-like nature; upon aging, however, a subset of proteins indeed shows a certain extent of viscoelastic transition (partial solidification) (Fig. 3). We found that gradual solidification does not mandatorily corroborate with amyloid fibril formation. Solid-like transition of liquid condensate might also occur due to crystal-like packing/ amorphous aggregation in the dense LLPS milieu. This suggests that solidification of liquid condensates might be specific to proteins in respect to sequence/structure and also could preserve the structure (therefore protein function) of most of the proteins^1,2,6,22,57^.

Although *C*_*LLPS*_ for most of the proteins is highly dependent on their charge (Fig. 4a), suggesting electrostatic interactions play a role in LLPS, a subset of proteins undergo LLPS either with exposed hydrophobic surfaces or in a combination of hydrophobic/electrostatic/H-bonding interactions. In this context, recent studies suggest that hydrophobic interaction can also play a significant role in the LLPS of proteins^18,61-63^. LLPS is also dependent on the solubility of the proteins, which strongly correlates with their intermolecular interaction strength. It is also dictated by the molecular weight (polymer length/amino acid number in proteins) and the nature of amino acid side chains^6,11,29,35,69^. For example, a stretch of a glycine-rich polypeptide with higher polypeptide flexibility and the absence of sidechain polyvalency might decrease the extent of intermolecular interaction^26^. However, hydrophobic amino acid (Val) and other aromatic amino acids might increase the interaction strength due to hydrophobic and other interactions (such as cation-π)^23,26,30^ when present in proteins. This interaction strength is highly reflected in *C*_*LLPS*_ as (Gly)_10_ requires double the polypeptide concentration (2 mM) for LLPS in comparison to (Val)_10_ (1 mM). Further, homopolymers of charged amino acids might not undergo LLPS due to repulsion unless their charges are neutralized. However, the segregation of oppositely charged residue in proteins is known to increase the tendency of phase separation^66^. Indeed, (Arg)_10_ and (Asp)_10_ homopolymers undergo LLPS either in presence of salt^24,39,46,73,74^ (Fig. 5b) or when they are mixed together (Fig. 5g). Another interesting observation is that different protein combinations undergo co-LLPS when mixed suggesting that nonspecific interactions might be responsible for heterotypic phase separation (co-LLPS). It is important to note that the requirement of a specific ratios of concentrations (such as *C*_*LLPS*_ for both components) as well as protein combinations might play a significant role in heterotypic phase separation.

However, if all proteins have the propensity for LLPS then how do cells maintain the native and/or soluble form of proteins? This is because the cellular protein concentrations mostly reside below their solubility limit^35^. Further, the cells might maintain the protein solubility by modifying the protein milieu; for example, by introducing post-translational modifications (phosphorylation, acetylation, etc.)^1,30,75,76^; or by other cellular factors such as ATP (natural hydrotrope)^38^, which is known to disrupt LLPS of a wide range of proteins. Moreover, proteins with highly charged surfaces might not be able to self-assemble due to *in-cell* ionic strength/salt concentration.

LLPS might also be tightly regulated based on the protein localization in specific organelles where it performs its native function,^1,5,22,45,77,78^ and the presence of DNA/RNA or other co-factors in cell^24,45-48,73,74,79,80^.

In conclusion, our study suggests that proteins/polypeptides with different structures and sequences can undergo LLPS although with different *C*_*LLPS*_. The presence of IDRs might provide an advantage in undergoing phase separation as they have higher polyvalency as well as a low structural order, resulting in substantially a greater number of molecular interactions^1,6,14,18,23^. Moreover, once a protein undergoes LLPS, its subsequent solidification might require very high concentration and/or specific interactions. Deregulation of protein quality control and turnover mechanisms in cells might pave the way for aberrant phase transition^1,6,12,81^. The similar generic state hypothesis has also been proposed for amyloid fibrils^82^ by proteins and polypeptides with an argument that cellular and subcellular conditions, protein quality control machinery and protein expression/post-translational modification do not allow such transition in cells. Also, Nature perhaps has evolved with a ‘negative design’ for proteins, which prevents amyloidogenesis^83^. Similar to this, we suggest that proteins can exist in multiple states in the cellular milieu. If LLPS is helpful for cellular fitness, cells might actively regulate their concentration and other factors favoring LLPS and vice versa (Fig. S22). However, a significant imbalance in this regulation can result in aberrant phase transition causing cell death, which eventually can lead to disease pathogenesis over time.^1,2,6,17,18,75^

## Methods

The *in vitro* LLPS of all proteins (commercially purchased, expressed/purified) and polypeptides was performed in presence of 20 mM sodium phosphate buffer (pH 7.4) and varying concentrations of PEG-8000. The condensate formation was examined under fluorescence microscopy. The liquid-like nature of the condensates was characterised through FRAP, fusion and temperature reversibility experiments. Liquid-to-solid transition was monitored using fluorescence recovery after photobleaching after 48 h of liquid condensate formation. The secondary structural transition of the various proteins immediately after LLPS (0 h) and 48 h of incubation were performed using CD and FTIR spectroscopy. Protein exposed hydrophobic surface and amyloid fibril formation during LLPS and liquid to solid-like transition were done using ANS and ThT fluorescence spectroscopy, respectively. The morphology of the condensates was analysed using TEM. The electrostatic and hydrophobic nature of proteins governing LLPS was studied using light scattering measurements (350 nm) in presence of NaCl and 1,6-hexanediol, respectively. Protein solubility was determined through ammonium sulfate precipitation study. Further, multicomponent LLPS by various combinations of proteins were performed using different proteins of various ratios and studied under confocal microscopy. The detailed methods are provided in Supplementary Methods.

## Supplementary Information

The supplementary information file contains materials and methods, Supplementary Fig. 1-22, Supplementary Tables 1-3 and 19 references.

## Author Contributions

† M.P. and K.P. contributed equally to this work. M.P., K.P., A.S.S., L.G., P.K., S.M., S.Maiti., D.C., R.B., and N.G. performed the *in vitro* experiments. M.P., K.P., D.D., S.R., and A.N., participated in analyzing the data. The study was conceived by S.K.M. and designed by S.K.M., R.P., M.P., and K.P. P.K. prepared the illustration. All authors participated in the writing of the manuscript and approved the final version.

## Data availability statement

The authors declare that all the data supporting the findings of this study are available within the paper and in supplementary information files. All the data analysis was performed using published tools and packages and has been cited in the paper and supplementary information text.

## ACKNOWLEDGMENT

We acknowledge IIT Bombay central facilities and SAIF for TEM, FTIR, LC-MS, MALDI and Confocal microscopy facility. We thank Prof. Sudipta Maiti, TIFR, Mumbai for helping us to utilize the HPLC facility. The authors acknowledge TATA Innovation (BT/HRD/35/01/03/2020), DST-SERB (File no. CRG/2019/001133), and DBT-Basic Science (File no. BT/PR22749/BRB/10/1576/2016) for financial support. M.P. acknowledges DST-INSPIRE, Government of India for the fellowship.

