## Supplementary Information for "Liquid condensate is a common state of proteins and polypeptides at the regime of high intermolecular interactions"

**Contents** **Page no.**


1. Materials S3
2. Supplementary Methods S3 – S15
3. Supplementary Figures (1 - 22) S16 – S37
4. Supplementary Table (1 – 3) S38 – S41
5. References S42

### Materials

All the reagents and chemicals used for the study were purchased from Sigma (USA) unless mentioned otherwise. The product information of the proteins is provided in Supplementary Tables 1 and 2. NHS-Rhodamine (Catalog no. 46406) and Fluorescein-5 isothiocyanate (FITC) (Catalog no. F1906) were procured from ThermoFisher Scientific (USA). Protease inhibitor cocktail (PIC) was obtained from Roche Applied Science (Catalog no. 05056489001). 1-Hydroxybenzotriazole hydrate (HOBt) (Catalog no. 157260), Triisopropylsilane (TIPS) (Catalog no. 233781), Trifluoroacetic acid (TFA) (Catalog no. T6508) and *N*, *N*'-Diisopropylcarbodiimide (DIC) (Catalog no. D4781) were purchased from Sigma (USA). *N*, *N*-Dimethylformamide (DMF) (Catalog no. 8.22275.2521), Dichloromethane (DCM) (Catalog no. 1.94508.2521), Acetonitrile (ACN) (Catalog no. 60003025001730), and Diethyl ether (Catalog no. 1.07026.0521) were purchased from Merck Millipore. Wang resin (100–200 mesh, 0.7 mmol/ g) (Catalog no. 8.55002) and 4-(Dimethylamino)pyridine (DMAP) (Catalog no. 8.51055) were purchased from Novabiochem (Germany). The polypeptides, pentaglycine (Catalog no. G5755), and hexaglycine (Catalog no. G5630) were purchased from Sigma-Aldrich (USA).

### Methods

#### Expression and purification of α-synuclein (α-Syn) and Tau protein

α-Syn was expressed and purified using previously established protocols with slight modifications[1,2]. Briefly, competent *E. coli* BL21 (DE3) cells were transformed using cloned plasmid and the expression was induced using isopropyl-β-D-thiogalactoside (IPTG) (1 mM). Following which the cells were centrifuged at 4000 rpm for 30 min at 4 °C. The pellet was resuspended in lysis buffer (50 mM Tris, 10 mM EDTA, 150 mM NaCl) and PIC (Roche) was added to prevent proteolytic cleavage. The cells were further lysed using a probe sonicator

(Sonics & Materials Inc.) at 40% amplitude with 3 s ON and 1 s OFF pulse for 10 min. The solution was then heated at 95 °C for 20 min and centrifuged at 9000 rpm for 30 min. The supernatant was used for nucleic acid precipitation using 10% streptomycin sulfate (136 μl/ml) and glacial acetic acid (228 μl/ml). The solution was then centrifuged at 9000 rpm for 30 min at 4 °C to remove nucleic acid. Following this, the protein precipitation was carried out using saturated ammonium sulfate (equal volume). The solution was kept at 4 °C for 4 h for complete precipitation and centrifuged at 10,000 rpm for 30 min at 4 °C. The protein was further washed using ammonium sulfate solution (50%) and centrifuged at 10,000 rpm. Finally, the protein was washed using ammonium acetate (100 mM) and precipitated using ethanol. This step was repeated three times. The solution was centrifuged and the pellet was dissolved in a minimum volume of ammonium acetate (100 mM) and lyophilized. The lyophilized protein was redissolved in 20 mM sodium phosphate buffer, pH 7.4, and further purified using size exclusion chromatography (SEC) in the Q Sepharose column before the LLPS experiment. The purity of the protein was confirmed by SDS-PAGE and Coomassie blue staining method.

Expression of full-length wild-type Tau protein (2N4R isoform containing 441 residues) was carried out by transforming tau/pET29b plasmid (Addgene id 16316) into *E. coli* BL21 (DE3) competent cells. The expression and purification protocol of Tau protein were similar to α-Syn with minor modification. Briefly, bacterial cells were grown in the presence of Kanamycin in Luria broth (LB) media at 37 °C to an optical density value between 0.7-1. Protein expression was induced with 1 mM IPTG followed by 4 h incubation at 37 °C in 200 rpm rotation. Cells were harvested by centrifugation and resuspended in 60 ml of lysis buffer (50 mM Tris, 10 mM EDTA, and 150 mM NaCl, 5 mM DTT at pH 8.0). PIC was added to the lysis buffer to prevent proteolytic cleavage. The cells were lysed by sonication (40% amplitude, 3 s ON and 1 s OFF) for 15 min using a probe sonicator (Sonics and Materials Inc., USA) and heat-denatured in hot water at 95 °C for 20 min. Cell debris and other denatured proteins were pelleted down by centrifugation at 10,000 rpm, 4 °C for 30 min. DNA was precipitated from the supernatant using streptomycin sulfate [10% (w/w)] and glacial acetic acid. After DNA removal, an equal volume of saturated ammonium sulfate was added and incubated at 4 °C overnight for protein precipitation. The solution was centrifuged twice at 12,000 rpm, 4 °C for 30 min. Pellet was dissolved in 100 mM ammonium acetate and reprecipitated in an equal volume of ethanol. The final pellet was redissolved in a minimum volume of 100 mM ammonium acetate, flash-frozen with liquid nitrogen, and lyophilized. The lyophilized protein powder was stored at -20 °C until used for experiments. The required amount of protein was dissolved in equilibrating buffer (20 mM sodium phosphate buffer, 1mM DTT) and further purified by size exclusion chromatography in the Q Sepharose column before the experiment. The purity of the protein was confirmed by the standard SDS-PAGE and Coomassie blue staining method.

**Size exclusion chromatography (SEC) of all proteins**.

All the commercially purchased proteins and recombinantly expressed/purified α-Syn and Tau were dissolved in a filtered 20 mM sodium phosphate buffer (pH 7.4, 0.01% sodium azide). The Superdex 200 TM 10/300 SEC column was pre-equilibrated with 3 column volumes of 20 mM sodium phosphate buffer (pH 7.4, 0.01% sodium azide) and the protein solutions were injected into the column. The proteins were isolated and the purity of the protein from SEC was confirmed using SDS-PAGE. The protein concentrations were determined by Beer-Lamberts law (c= $A/\varepsilon l$), where $c$ is the protein concentration in molar, $l$ is the path length in cm, $A$ is the absorbance value at respective wavelength, and $\varepsilon$ is the molar absorption coefficient at the respective wavelength, using UV spectroscopy (Jasco V650, Japan). The absorbance measurement at 280 ($A_{280}$) was used for determining the protein concentration for all the proteins except the chromophore-containing proteins such as Hb ($A_{406}$, $\varepsilon_{406}$=270548 $M^{-1}cm^{-1}$)[3], Mb ($A_{408}$, $\varepsilon_{408}$=129000 $M^{-1}cm^{-1}$)[4], Cyt c ($A_{410}$, $\varepsilon_{410}$=101600 $M^{-1}cm^{-1}$)[5], and CATA

($A_{405}$, $\varepsilon_{405}$=324000 $M^{-1}cm^{-1}$)[6] whose protein concentration was determined using the extinction coefficient of the respective chromophore group.

**Solid-phase peptide synthesis.**

All the peptides were synthesized by 9-fluorenylmethoxy- carbonyl (FMOC) chemistry using the manual solid-phase peptide synthesis method[7]. The synthesis was performed with the scale of 0.20–0.25 mmol on a Wang resin. In a typical synthesis, the first amino acid was loaded on Wang resin by dissolving 1 eq. of amino acid and HOBt in DMF, followed by the addition of 1 eq. of DIC and finally DMAP in catalytic amt. (0.1 eq.). The coupling was kept for 2-3 h and washed several times with DMF and DCM after completion of the reaction. The FMOC group was removed using 25% piperidine in DMF. The next coupling was repeated using DIC/HOBt coupling agent with the equivalent amount of the next amino acid. After the synthesis of the desired length polypeptide, the peptide was cleaved off from the resin using a standard cleavage cocktail, TFA: Phenol: TIPS: water (88:5:2:5). Further, the cleavage solution was transferred into an ice-cold ether solution to get the precipitated peptides. After precipitation, the ether solution was evaporated and the peptides were redissolved in ammonium bicarbonate (50 mM).

The synthesized peptides [$(Gly)_{10}$, $(Asp)_{10}$, and $(Arg)_{10}$] were purified using HPLC equipped with a reverse phase-C18 column. The mobile phase was used with the 90 min gradient system starting from 10% ACN/water (0.1% TFA) to 90% ACN/water system with a flow rate of 1 ml/min. The samples were injected from a 5 mg/ml stock concentration and 200 μl of peptide aliquot solution was injected using an autosampler injector. The instrument was provided with a UV-Vis detector (dual-wavelength) and absorbance at 195 nm was recorded. For analysis we used the data acquired at 195 nm (the analytes had maximum molar absorptivity). Using these parameters, all the synthesized polypeptides were separated. However, $(Val)_{10}$ was not purified using HPLC, since it exhibits poor solubility in the given HPLC mobile phase gradient system and therefore, was used as synthesized. The purified polypeptides were characterized using ESI LC-MS and MALDI analysis.

**Fluorescent labeling of protein/peptide**.

The NHS-Rhodamine and FITC labeling of protein was done as per the manufacturer's protocol (ThermoFisher Scientific, USA). Briefly, 5X molar excess of FITC/rhodamine (dissolved in DMSO) was added to protein obtained after SEC. For FITC, the mixture was incubated on a magnetic stirrer at 4 °C for 6 h in dark with slow rotation. For NHS-Rhodamine labeling, the protein mixture was incubated for 2 h at room temperature in dark with slow stirring. The excess dye was removed by dialysis in 20 mM sodium phosphate buffer (pH 7.4) at 4 °C for 48 h, with regular buffer exchange in 6 h intervals. The concentration of the labeled protein was determined as per the manufacturer's protocol. The polypeptides were labeled as mentioned previously. The excess FITC/rhodamine dye was removed by dialysis in 20 mM sodium phosphate buffer (pH 7.4) for 12 h with regular buffer exchange in 2 h intervals at 4 °C. After dialysis, the labeled polypeptide solution was lyophilized and the concentration was determined by redissolving the dry weight in 20 mM sodium phosphate buffer (pH 7.4). For all experiments, we used 1:10 (v/v) of labeled versus unlabeled protein/polypeptide, unless mentioned otherwise.

***In vitro* liquid-liquid phase separation of proteins**.

For LLPS experiments, acid-treated coverslips were used[8]. To do this, the glass slides and 12 mm coverslips (Blue Star, India) were kept in aqua regia [1:3 (v/v) nitric acid/hydrochloric acid] for 12 h and thoroughly washed with Milli-Q. After every wash, the pH of the Milli-Q was checked until it reached 7.0. The slides and coverslips were air-dried in a laminar air-flow hood under sterile conditions and used for all the subsequent LLPS experiments.

For LLPS, the proteins after SEC were used to prepare the reaction mixture at different protein and PEG-8000 concentration [(0%, 5% 10%, 15% and 20% (w/v)] in 20 mM sodium phosphate buffer (pH 7.4, 0.01% sodium azide) to determine the phase regime. The polypeptides $(Arg)_{10}$ and $(Asp)_{10}$ were dissolved in 20 mM sodium phosphate buffer (pH 7.4). For $(Val)_{10}$ and $(Gly)_{10}$, 1 mg of the respective polypeptide was dissolved in 20 µl of TFA to obtain a homogenous solution and the volume was adjusted to 50 µl by addition of 20 mM sodium phosphate buffer (pH 7.4). TFA was removed by nitrogen gas purging and 20 mM sodium phosphate buffer was added to obtain a stock solution of 5 mM for both the polypeptides. For LLPS of polypeptides, the reaction mixture at different polypeptide concentrations and PEG-8000 concentrations [(0%, 5% 10%, 15% and 20% (w/v)] in 20 mM sodium phosphate buffer (pH 7.4, 0.01% sodium azide) was prepared to determine the phase regime. The reaction mixture (proteins/polypeptides) was drop-casted on the acid-treated slides and sandwiched with an acid-treated 12 mm glass coverslip (Blue Star, India). The coverslips were sealed using commercially available nail paint. The slides were incubated at 37 °C in a moist chamber and phase separation was monitored using 63X oil immersion objective in the DIC (Differential Interference contrast) mode and fluorescence mode under a DMi8 microscope (Leica Microsystems, Germany). All the images were obtained at 16-bit depth with 2048 x 2048 pixels resolution unless mentioned otherwise. The images were analyzed using ImageJ (NIH, Bethesda, USA) software.

**Fluorescence and confocal microscopy.**

The *in vitro* LLPS and liquid condensate formation for all the NHS-Rhodamine labeled proteins [1:10 (v/v) labeled to unlabeled protein] were observed using a DMi8 microscope (Leica Microsystems, Germany) under DIC and fluorescence mode using an appropriate fluorescence channel (560 nm) at 16-bit depth with 2048 x 2048 pixels resolution. The FRAP analysis at $C_{LLPS}$ of proteins (0 h and 48 h after LLPS) and the heterotypic co-LLPS studies were performed using a laser scanning confocal microscope (LSM 780 Zeiss Axio-Observer Z1 microscope (inverted)) equipped with iPlan-apochromat 63X/1.4 NA oil immersion objective and with appropriate fluorescence channel (488 nm and 560 nm). The images were obtained with a frame size of 1024 pixels X 1024 pixels with 8 bit-depth unless mentioned otherwise. The images were processed using ImageJ (NIH, Bethesda, USA) software.

**Light scattering measurements**.

The $C_{LLPS}$ for LLPS of the respective protein sample in presence of 10% PEG-8000 (w/v) (LLPS-inducing condition) was used for static light scattering measurements. The excitation and emission wavelength were set at 350 nm and the slit width was kept at 5 nm for both. The measurements were acquired in continuous mode using a spectrofluorimeter (JASCO FP 8500, USA). The experiment was performed twice. For studying the effect of salt (NaCl) (disrupting electrostatic interaction) and 1,6-hexanediol (disrupting hydrophobic interaction), the protein samples at their respective $C_{LLPS}$ in the presence of 10% (w/v) PEG-8000 and NaCl (150 mM) or 1,6-hexanediol (10%) (w/v) was used for the measurements. The experiment was performed twice. A plot of light scattering intensity against time resulted in a sigmoidal curve and the data were background corrected, normalized, and fitted using the Boltzmann equation and $t_{1/2}$ was calculated as follows;

$$y = y_0 + (y_{\max} - y_0) / [1 + e^{-(k(t-t_{1/2}))}] \qquad (1)$$

Where, $y$ = the light scattering intensity at a particular time point, $y_{max}$ = maximum light scattering intensity, $y_0$ = light scattering values at $t_0$. The data was plotted using OriginPro 2021 (Origin Lab, USA) software.

**Fluorescence Recovery After Photobleaching (FRAP).**

For FRAP experiments, NHS-rhodamine labeled [10% labeled and 90% unlabeled (v/v)] protein mixture in presence of PEG-8000 (10% w/v) at respective $C_{LLPS}$ were incubated in eppendorf at 37 °C in a moist chamber for LLPS. At different time points (0 h and 48 h) the samples were drop-casted on acid-treated glass slides and covered with 12 mm acid-treated coverslip. The condensate was bleached and fluorescence recovery was determined using a previously established protocol[9]. The experiments were performed using a built-in FRAP module in Zeiss Axio-Observer Z1 confocal microscope with 63X oil-immersion objective (NA 1.4). A 561 nm DPSS 561-10 laser (at 100% laser power) was used to bleach the center of the condensate and two other regions of interest (ROI) with the same diameter were also recorded to determine the background and passive bleaching corrections. The fluorescence intensity after bleaching was simultaneously recorded for all three ROIs using the Zen Pro 2011 (Zeiss, Germany) software provided with the instrument. The images were obtained with a frame size of 512 pixels X 512 pixels with 8 bit-depth. For the heterotypic LLPS experiment, two ROIs were bleached using 561 nm DPSS 561-10 laser and 488 laser (30% and 100% laser power, respectively), and the fluorescent intensity was measured for all the ROIs with respective background and bleach correction for an individual channel. The diameter of all the ROIs was kept identical. The images were acquired with the frame size of 512 X 512 pixels with 8 bit-depth. According to the previously published protocol[9], the fluorescence recovery data was background corrected, normalized, and fitted using the single exponential recovery function in OriginPro 2021 (Origin Lab, USA) software, and $t_{1/2}$ was determined. The equation used for fitting is as follows[10-14];

$$I\,(t) = A\left(1 - exp\,\left(\frac{-t}{\tau}\right)\right) + C \qquad (2)$$

Where, τ is the fluorescence recovery time constant, 'A' corresponds to the mobile fraction of the fluorescent probe, and '$C$' is the Y-intercept of the recovery curve.

The half time of the recovery ($t_{1/2}$) was calculated from,

$$t_{1/2} = \tau\, ln(2) \qquad (3)$$

**Thioflavin T (ThT) fluorescence assay**.

For ThT fluorescence assay, 100 µl of unlabeled SEC isolated protein samples (proteins which showed low recovery of fluorescence at 48 h after LLPS; β-cas, CATA, GG, LT, Tau, and α-Syn) was diluted in 20 mM sodium phosphate buffer (pH 7.4, 0.01% sodium azide) at a final concentration of 10 µM in presence of PEG-8000 (10% w/v) and incubated in 37 °C for LLPS (0 h and 48 h). At both time points, ThT fluorescence measurements were performed. To do that 1 µl of 1mM ThT dye (Tris-HCl buffer, pH 8.0, 0.01% sodium azide) was added to the protein samples. ThT fluorescence measurements were recorded using Spectrofluorimeter (JASCO FP 8500, USA) instrument at an excitation wavelength of 450 nm and an emission range of 460-500 nm with the slit width of 5 nm for both excitation and emission measurement. The graph was plotted using the OriginPro 2021 (Origin Lab, USA) software at the emission maxima ($\lambda_{max}$~ 480 nm) after background corrections. The experiment was repeated two times.

**Transmission Electron Microscopy (TEM)**.

TEM analysis was performed for a subset of proteins (proteins that showed substantial solidification using FRAP data; α-Syn, LT, GG, Tau, β-cas, and CATA) at their respective $C_{LLPS}$ immediately after LLPS (0 h) and after 48 h of incubation. For sample preparation, the coverslip containing LLPS solution was removed from the slide of LLPS samples (0 h and 48 h) and it was directly transferred on the EM grid (Electron Microscopy Sciences, USA). The grids were stained using uranyl formate (1% v/v) for 5 min and excess dye was removed with the help of filter paper. The grids were directly air-dried without any further washes before imaging. Imaging was done using transmission electron microscopy (TEM, JEOL JEM 2100F, Japan) at 200 kV with 10000X magnification. The images were recorded digitally using Gatan Soft imaging system (Japan).

**8-anilino-1-naphthalenesulfonic acid (ANS) binding assay**.

To determine the extent of the exposed hydrophobic surface of proteins, ANS fluorescence binding assay was performed. Briefly, 3 µl of 5 mM ANS was added to 100 µl of all the SEC protein samples in presence of PEG-8000 (10% w/v) in 20 mM sodium phosphate buffer (pH 7.4, 0.01% sodium azide) at the final concentration of 10 µM. The mixture was incubated for 5 min in dark at room temperature. The fluorescence intensity measurements were acquired using a spectrofluorimeter (JASCO FP 8500, USA) with 370 nm as an excitation wavelength and 400-600 nm as an emission wavelength range. The slit width was set to 5 nm for both excitation and emission wavelength. The acquired spectra were plotted after background correction using GraphPad Prism 8 at the emission wavelength of 475 nm. The experiment was repeated two times.

**Circular dichroism (CD) study**.

The far-UV circular dichroism spectra for SEC isolated proteins before LLPS (with PEG-8000 10% w/v, non-LLPS inducing condition), immediately after LLPS (0 h), and after 48 incubations were recorded using JASCO-1500 CD spectrophotometer (USA) in a 0.1 cm microcuvette (Hellma Forest Hills, NY). 200 µl of each protein sample was diluted to the final concentration of 10 µM in 20 mM sodium phosphate buffer (pH 7.4, 0.01% sodium azide) and used for CD measurement. The spectra were recorded for the wavelength range of 260-198 nm at 20 °C with a scanning speed of 200 nm/min. Three accumulations for each sample were acquired and the experiment was done in duplicate. The buffer subtraction and smoothing of the data were done as per the manufacturer's instructions. The data was plotted using KaleidaGraph software.

**Fourier-transform infrared (FTIR) spectroscopy.**

FTIR spectroscopy was performed to determine the secondary structure of the proteins. The SEC isolated proteins before LLPS (with PEG-8000 10% w/v, non-LLPS inducing condition), immediately after LLPS (0 h), and after 48 h incubation were spotted on the KBr pellet and were subsequently dried under IR (infra-red) lamp. 10 µl of each protein sample (GG, LT, CATA, Tau, α-Syn, and β-cas) at $C_{LLPS}$ was diluted to the final concentration of 10 µM in 20 mM sodium phosphate buffer (pH 7.4, 0.01% sodium azide) and used for FTIR measurement. Vertex 80 FTIR system equipped with a DTGS detector (Bruker, Leipzig, Germany) was used to record the spectra in the range of 1800–1500 $cm^{-1}$. Each spectrum was recorded using an average of 32 scans at a resolution of 4 $cm^{-1}$. Fourier self-deconvolution (FSD) method was used to deconvolute the spectra corresponding to the wavenumbers 1700−1600 $cm^{-1}$ [15]. The Lorentzian curve fitting procedure was employed to fit the spectra using Opus-65 software (Bruker, Leipzig, Germany) as per the manufacturer's instruction. The experiments were performed twice with similar observations.

**Protein sequence analysis and correlation plot parameters.**

The total number of amino acids, charged and aromatic residues were counted using Expasy ProtParam tool from the sequences obtained from Uniprot (Table S1). The quantity absolute charge, $\sqrt{(N^2+P^2)}$ (N = number of negative residues and P = number of positive residues) was calculated from the output values of ProtParam. Kappa (κ) values were estimated using CIDER [16] analysis tool online. κ is the measure of the extent of charge segregation along the sequence length, where a larger value indicates high segregation of positive and negative charges. The κ values of Hb and CATA were calculated as the average values of the respective protein chains (2 α and 2 β chains for Hb, and CATA was considered as tetramer). The κ value of Chymo was calculated considering a single sequence composed of chains A, B and C.

**Protein Solubility assay**.

To determine the relative solubility of all proteins, a solubility assay was performed using the ammonium sulfate precipitation method. To do that saturated ammonium sulfate solution (4 M) was prepared in deionized water. 120 µl of 20 µM SEC isolated protein samples in 20 mM sodium phosphate buffer (pH 7.4, 0.01% sodium azide) was titrated using increasing concentrations of ammonium sulfate. After each addition of ammonium sulfate, protein samples were incubated for 5 min and centrifuged at 12,000 rpm. The concentration of the supernatant was measured using UV spectroscopy (Jasco V650, Japan) at 280 nm except for the chromophore containing proteins (Hb, Mb, CATA, and Cyt c) whose concentration was measured as previously mentioned.

The concentration of each protein after every addition of ammonium sulfate was plotted against the concentration of the salt. The data were fitted using the Dose Response model after normalization. The $C_{1/2}$ value was calculated based on the concentration of salt at which 50% of the protein was precipitated. Since, LYS, CA and β-lac did not reach 50% precipitation we calculated the $C_{0.4}$ value which is the concentration of ammonium sulfate required to precipitate 40% of the protein. The data was plotted together for the purpose of relative correlation. All data were plotted and fitted using OriginPro 2021 (Origin Lab, USA) software.

**Heterotypic LLPS.**

For heterotypic LLPS, FITC labeled proteins were mixed with other NHS-Rhodamine labeled proteins in the presence of PEG-8000 (10% w/v) in 20 mM sodium phosphate buffer (pH 7.4, 0.01% sodium azide) at various protein concentrations (equal protein concentration, $C_{LLPS}$, ½ $C_{LLPS}$, and ¼ $C_{LLPS}$). For all the experiments, we used 1:10 (v/v) of labeled versus unlabeled protein, unless mentioned otherwise. The mixture was drop-casted on acid-treated glass slides and sandwiched with a 12 mm acid-treated coverslip. The coverslip was sealed using commercially available nail paint. The slides were incubated in a moist chamber at 37 $^{o}$C for 8 h and confocal microscopy (LSM 780 Zeiss Axio-Observer Z1 equipped with iPlan-apochromat 63X/1.4 NA oil immersion objective) was performed. The appropriate fluorescence channel (488 nm and 560 nm) was used for imaging. Notably, care was taken to prevent any bleed-through between FITC channel and Rhodamine channel by appropriately setting the emission wavelength range and gain settings for individual channel as control and similar settings were used during the acquisition of co-LLPS condensates. The images were obtained with a frame size of 1024 pixels X 1024 pixels with 8 bit-depth. The images were processed using ImageJ (NIH, Bethesda, USA) software.

**Supplementary Figures**

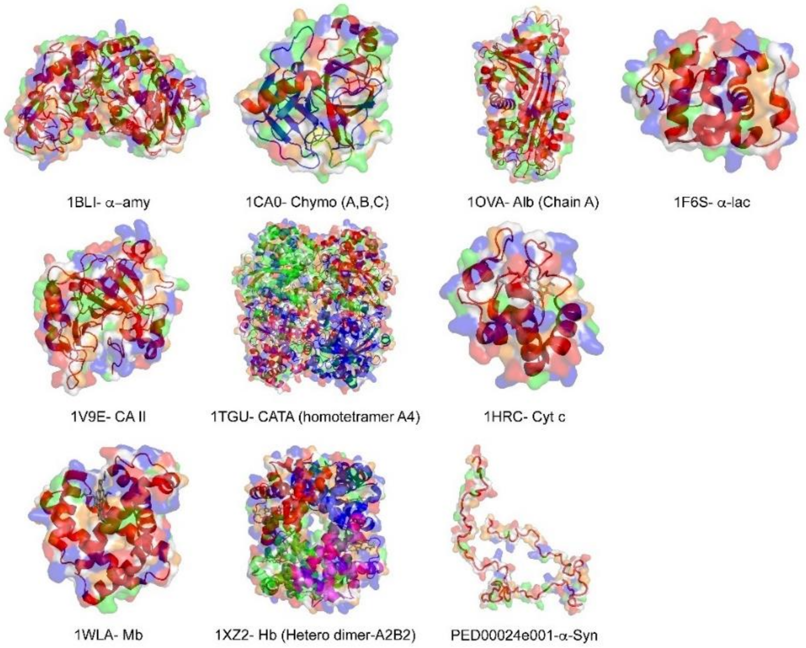


**Figure S1. The structure along with the surface plot of various proteins.** The structure of all the proteins was generated from their PDB identification code and represented with the surface plot using PYMOL. The data shows that all proteins exhibit different secondary structures and surface charge distribution. The color representations are red, negatively charged residues; blue, positively charged residues and green, hydrophobic residues.

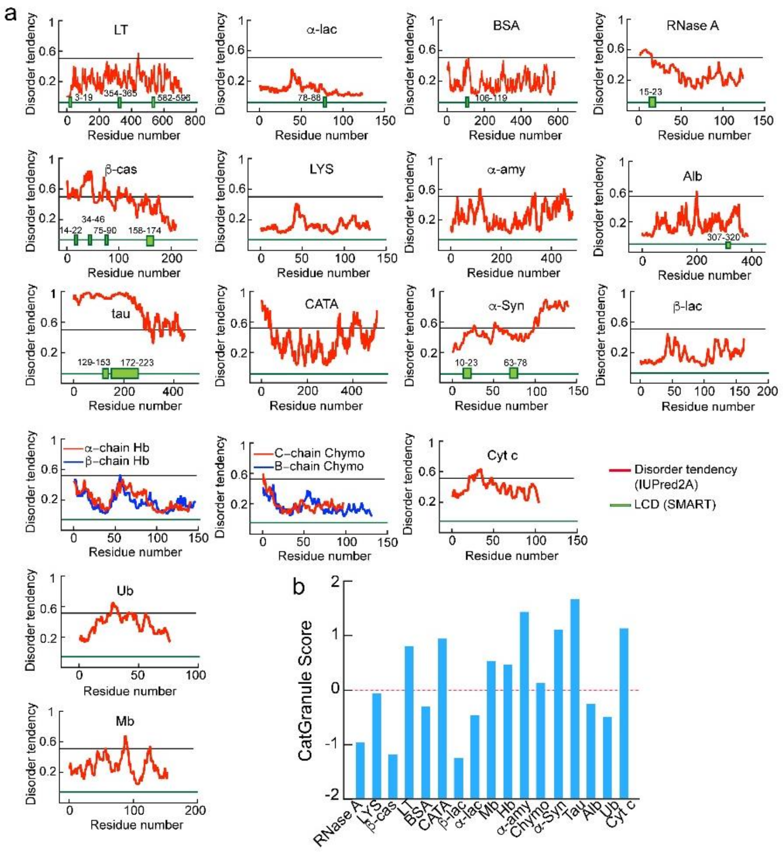

**Figure S2. *In silico* analysis of the primary sequence of the various protein.** *In silico* analysis of all the protein sequences showing (a) disorder tendency by IUPred2A[17], low complexity domains (LCDs) by SMART[18]. The disordered tendency is represented with red color where the threshold of 0.5 corresponds to the disordered region. The green color denotes the presence of LCDs. (b) propensity for LLPS determined using CatGranule[19]. The line indicates the threshold value for LLPS.

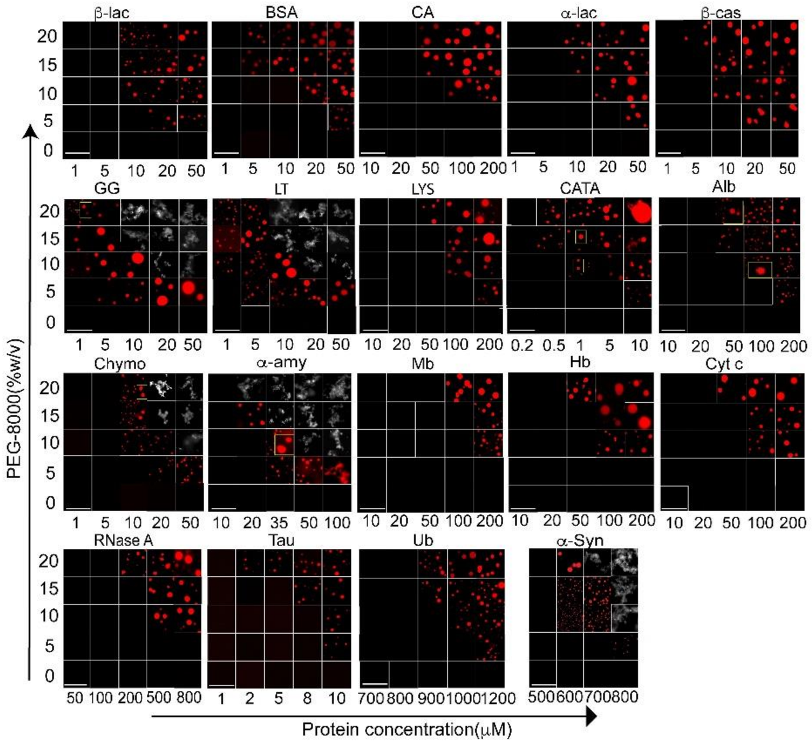


**Figure S3. The phase separation regime of different proteins.** The phase separation regime of all NHS-Rhodamine labeled proteins [1:10 (v/v) of labeled versus unlabeled protein] at varying PEG-8000 concentration (0%, 5%, 10%, 15% and 20% w/v) and protein concentrations in 20 mM sodium phosphate buffer at pH 7.4. The scale bar is 10 μm. The data showing low $C_{LLPS}$ for proteins such as LT and GG (1 μM) as well as very high $C_{LLPS}$ (≥500 μM) for proteins such as RNase A, Ub, and α-Syn. Representative images of precipitates are shown in 'grayscale' LUT. The experiment was repeated three times with similar observations.

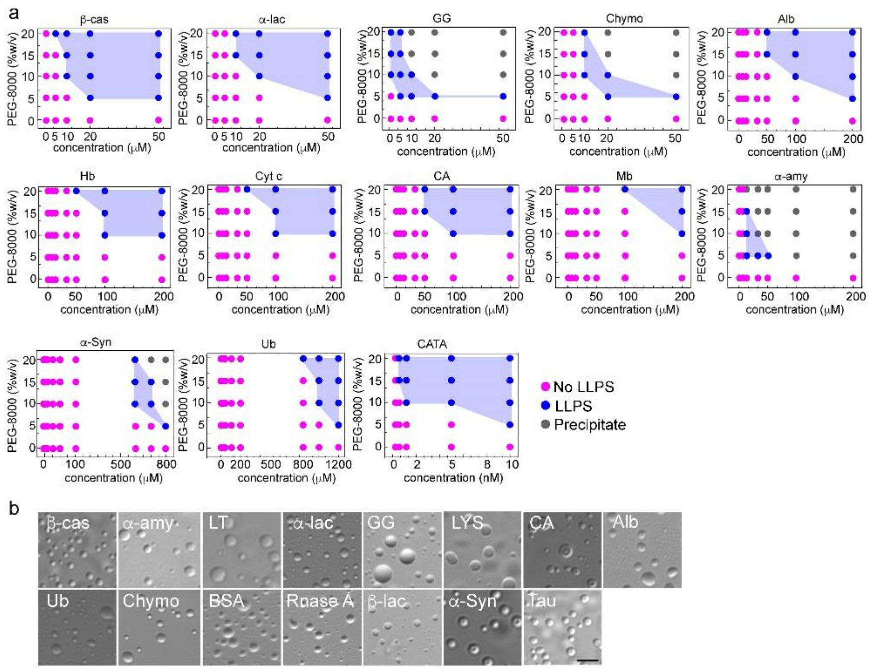


**Figure S4. Phase diagram of different proteins.** (a) The phase regime of all proteins depicting LLPS at varying protein and PEG-8000 concentrations (0%, 5%, 10%, 15% and 20% w/v) in 20 mM sodium phosphate buffer (pH 7.4). The different states are represented with various color codes. Pink color indicates no LLPS (soluble state), blue color indicated LLPS (condensate formation), and gray color indicates precipitates. (b) Representative DIC images of condensate formation for different proteins during light scattering measurements (350 nm) at their $C_{LLPS}$ in presence of 10% (w/v) PEG-8000 after nucleation. The scale bar is 5 µm.

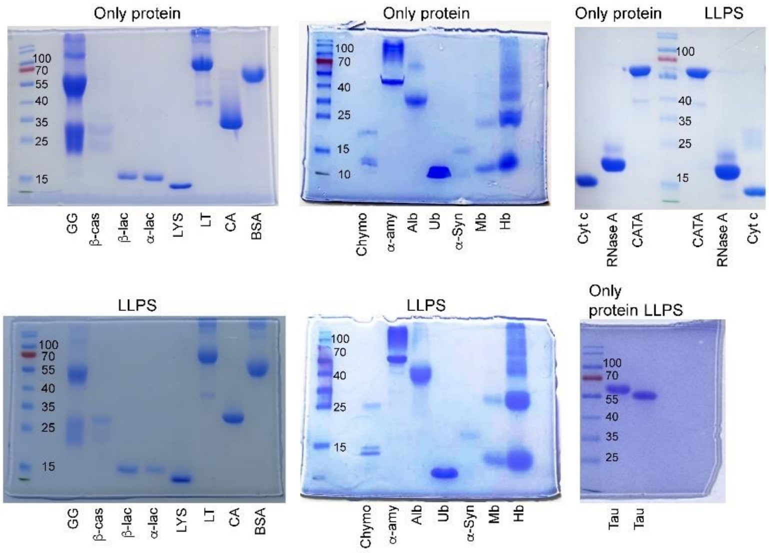


**Figure S5. SDS-PAGE of SEC purified proteins.** The SDS-PAGE images showing the presence of protein bands at their respective molecular weight and confirming the purity of the protein with no degradation even after LLPS.

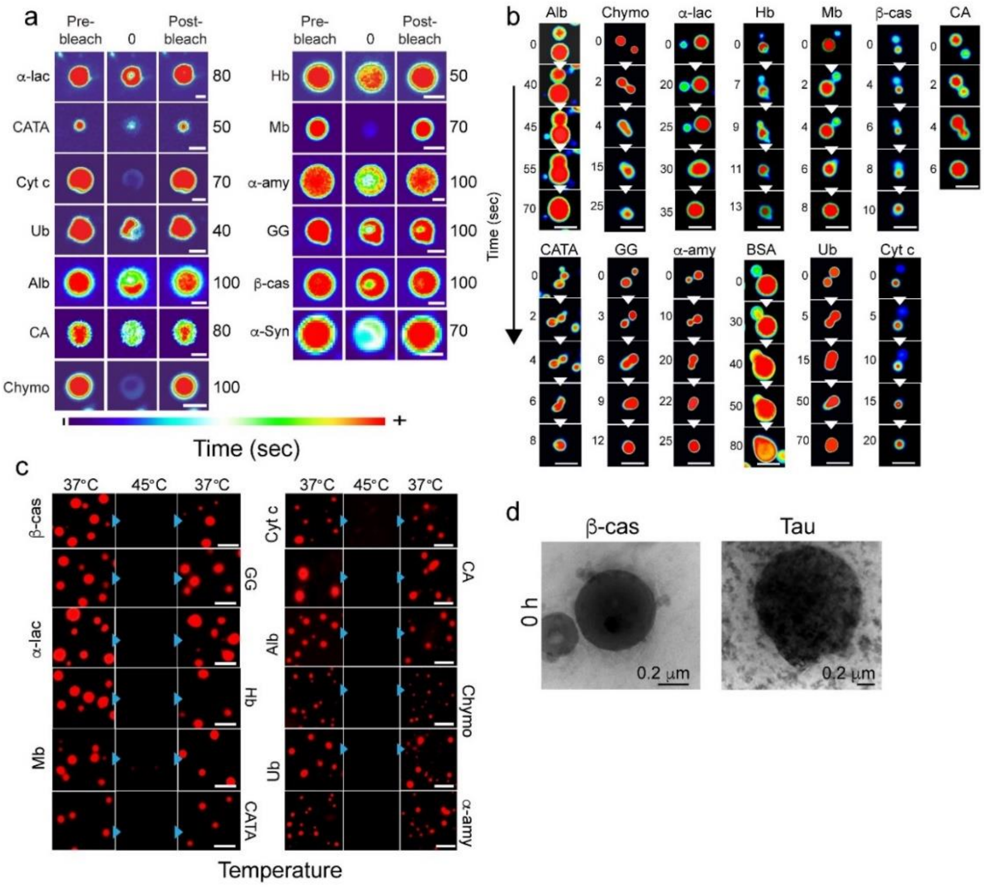


**Figure S6. Liquid-like nature of different protein condensates.** (a) Representative image showing the protein condensates immediately after LLPS (0 h) during FRAP analysis (before bleaching, at bleaching, and after bleaching). The images are represented in 'thermal' LUT for better visualization. The scale bar is 2 µm. (b) Fusion of liquid condensates. Representative time-lapse images of the proteins showing the fusion of small condensates upon contact, resulting in the formation of larger condensates over time. 'Royal' LUT was used for better visualization. The scale bar is 5 µm. (c) Thermo-reversibility of liquid condensates. The fluorescence microscopy images of the NHS-Rhodamine labeled [1/10 (v/v) labeled to unlabeled protein] condensates at their $C_{LLPS}$ in the presence of PEG-8000 (10% w/v) indicating thermo-reversibility (37 °C → 45 °C → 37 °C). The experiment was performed two times with similar observations. The scale bar is 5 µm. (d) Representative TEM images of β-cas and Tau showing the morphology of protein condensate at LLPS (0 h). *n=2* independent experiments were performed.

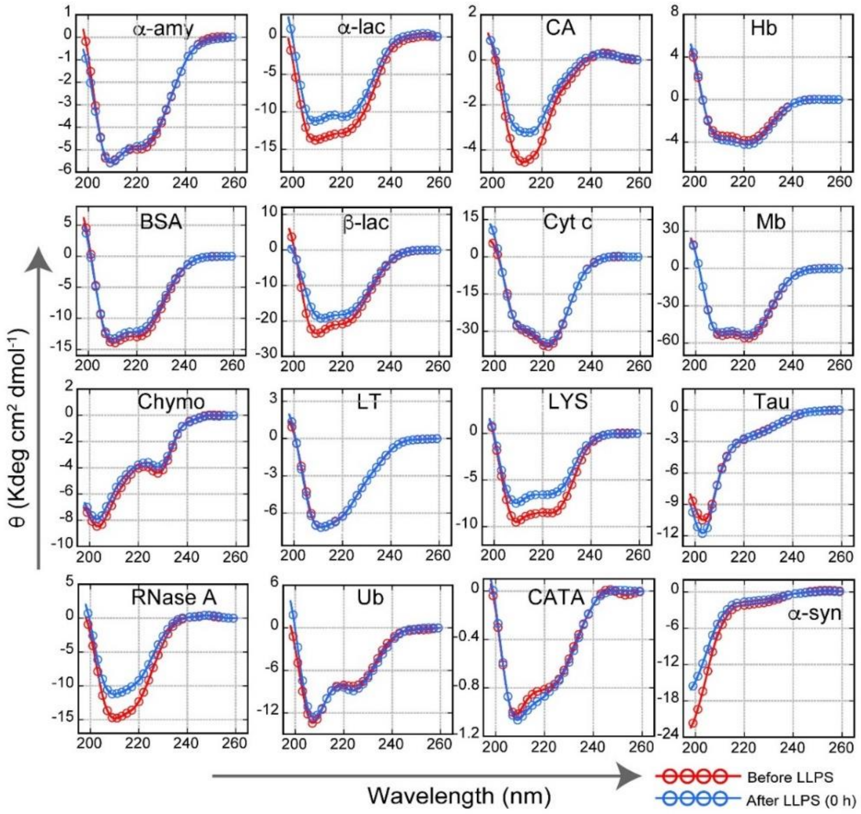


**Figure S7. Secondary structural analysis of various proteins.** The CD spectroscopic analysis of all proteins immediately after LLPS (0 h) demonstrates that most of the proteins did not undergo secondary structural transition due to LLPS. The red color represents the spectra of protein in presence of PEG-8000 (10% w/v) (without LLPS), and blue indicates CD spectra of protein immediately after LLPS (0 h) in 20 mM sodium phosphate buffer (pH 7.4) and 10% (w/v) PEG-8000.

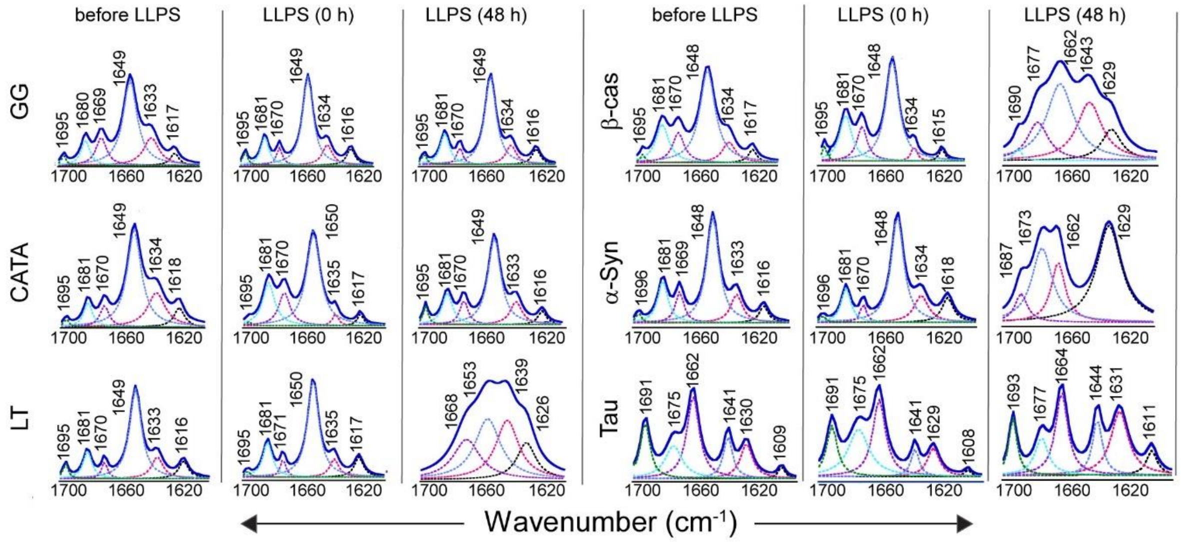


**Figure S8. Determination of the secondary structure of proteins using FTIR spectroscopy.** Deconvoluted FTIR spectra of GG, CATA, LT, Tau, α-Syn and β-cas showing the different secondary structures of protein immediately after addition of PEG-8000 (before LLPS) (**1st column**), immediately after phase separation (0 h) (**2nd column**), and after 48 h of incubation (**3rd column**). GG, CATA, and LT show a major peak at ~1650 $cm^{-1}$ which is characteristic of the α-helical structure, whereas β-cas and α-Syn shows a major peak at 1648 $cm^{-1}$, which is characteristic of random coil (RC) structure. The secondary structures of the proteins showing no significant change over different conditions through the LLPS time course except β-cas, α-Syn, and LT show a change in secondary structure over time. The experiment was performed two times with similar observations.

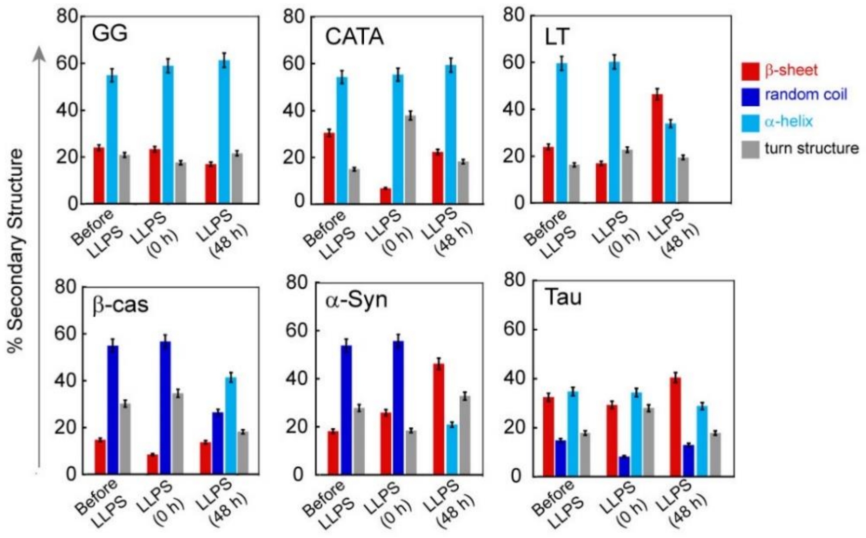


**Figure S9. Determination of the percentage of secondary structures by various proteins during LLPS and subsequent solidification.** The deconvoluted FTIR spectra were used for calculating the percentage of secondary structures. The percentage of secondary structures for GG, CATA, LT, Tau, α-Syn, and β-cas are shown at different conditions. The color representation is red, β-sheet conformation; dark blue, random coil; light blue, α-helix and gray, turn structure. The data values represent mean ± *s.d.* for *n=2* independent experiments.

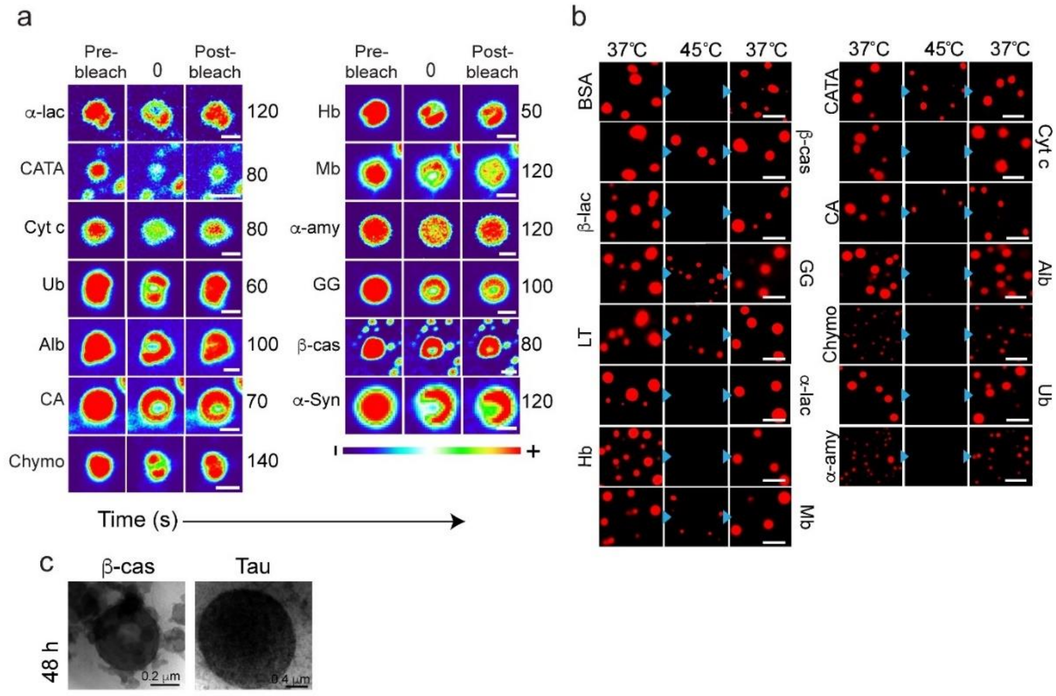


**Figure S10. The liquid-to-solid transition of various proteins.** (a) Representative image showing the liquid condensates at 48 h during FRAP analysis (before bleaching, at bleaching, and after bleaching). The data suggests the liquid-to-solid transition of the protein condensates. The images are represented in 'thermal' LUT for better visualization. The scale bar is 2 μm. (b) Thermo-reversibility of protein condensates. The fluorescence microscopy images of the NHS-Rhodamine labeled [1:10 (v/v) labeled to unlabeled protein] condensates at their $C_{LLPS}$ after 48 h of incubation in presence of PEG-8000 (10% w/v) indicating lack of thermo-reversibility (37 °C → 45 °C →37 °C) after ageing (48 h) for a subset of protein condensates. The experiment was performed two times with similar observations. The scale bar is 5 μm. (c) Representative TEM images of β-cas and Tau showing the morphology of protein condensate at LLPS (48 h). *n=2* independent experiments were performed.

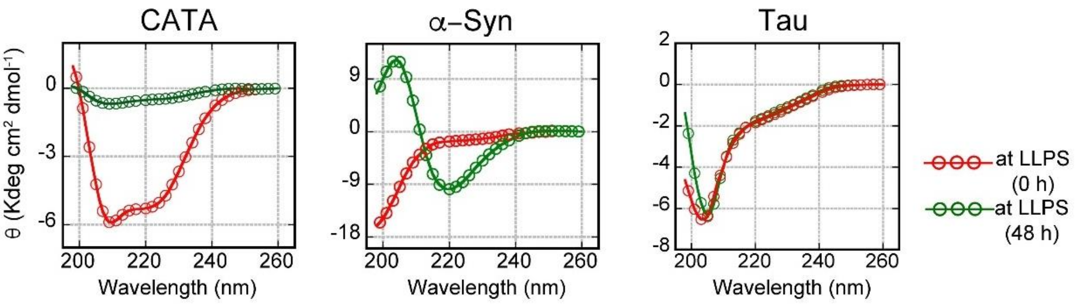


**Figure S11. Secondary structural analysis of various proteins after 48 h of LLPS.** The CD spectroscopic analysis of proteins (proteins that showed substantial solidification using FRAP data) after 48 h of LLPS showing a significant decrease in molar ellipticity (θ), indicating liquid-to-solid transition. The decrease in CD molar ellipticity however could be due to less protein solubility after ageing. The red color represents the CD spectra of protein immediately after LLPS (0 h) and the green represents the CD spectra of protein after 48 h of incubation. Protein LLPS was done in 20 mM sodium phosphate buffer (pH 7.4) in the presence of 10% (w/v) PEG-8000.

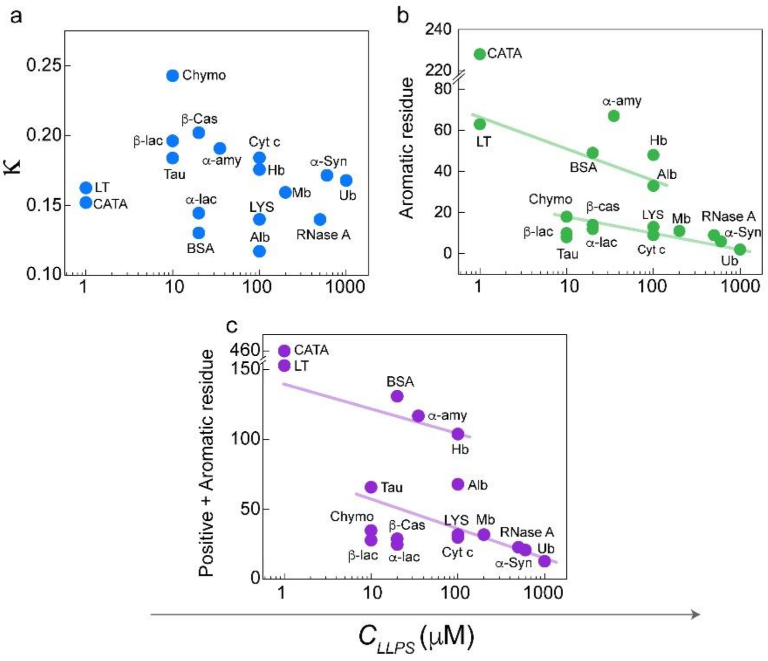


**Figure S12. Correlation plots of intrinsic parameters of the proteins with $C_{LLPS}$ (semi-log comparison)** (a) Kappa (κ) measurement was done using CIDER[16] to understand the extent of segregation of charges along the sequence length. High κ values indicate higher charge segregation along the length of the amino acid chain. (b) The distribution of the number of aromatic residues vs $C_{LLPS}$ showed two clusters of protein, showing negative linear correlation. Protein in one cluster (upper region) comprised majorly high MW proteins while those in the second cluster (lower region) comprised of majorly lower MW proteins. (c) Proteins with a higher number of positive and aromatic residues (a measure of cation-π interactions) showed a higher propensity for LLPS. Trend lines drawn (b, and c) are qualitative for guide purposes only.

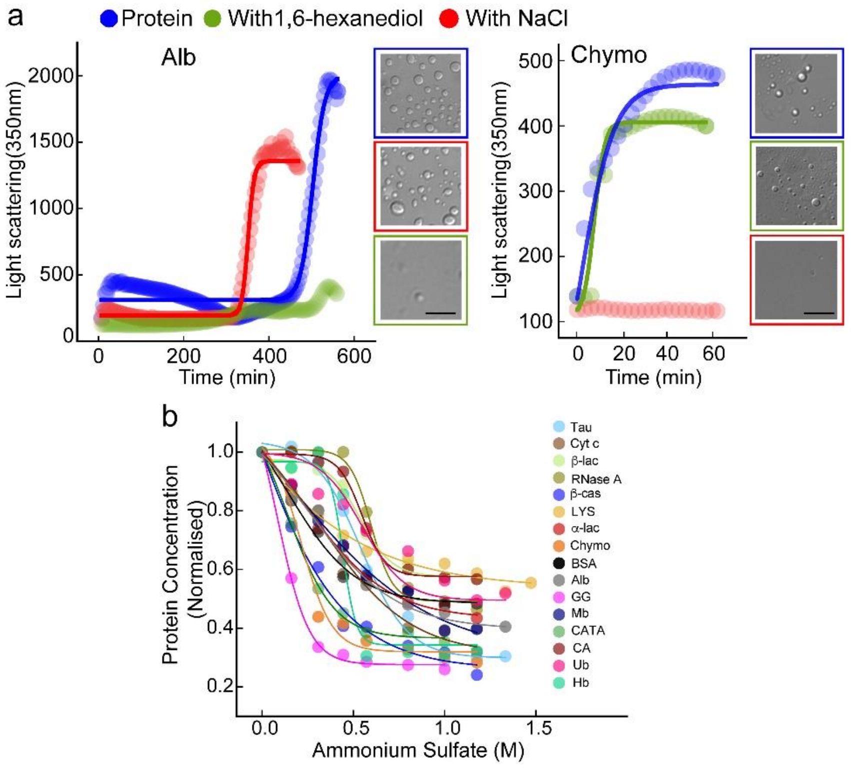


**Figure S13. Intermolecular interactions are responsible for protein LLPS.** (a) The LLPS study in the presence and absence of NaCl (150 mM) and 10% (w/v) 1,6-hexanediol to disrupt electrostatic and hydrophobic interactions, respectively. The static light scattering measurement at 350 nm of Alb and Chymo at their respective $C_{LLPS}$ in presence of 10% (w/v) PEG-8000 shows scattering due to LLPS. The color blue indicates spectra of protein in the presence of 10% (w/v) PEG-8000. Green and red colors denote the light scattering measurement of respective protein in presence of 1,6-hexanediol (10%) and NaCl (150 mM), respectively. Representative DIC images showing the absence or presence of condensate formation. The scale bar is 5 µm, *n=2,* independent experiments. (b) The decrease in protein concentration is due to the salting-out effect of various proteins using an increasing concentration of ammonium sulfate (0 to 4 M). The data represent the mean ± *s.d.* for *n=2* independent experiments.

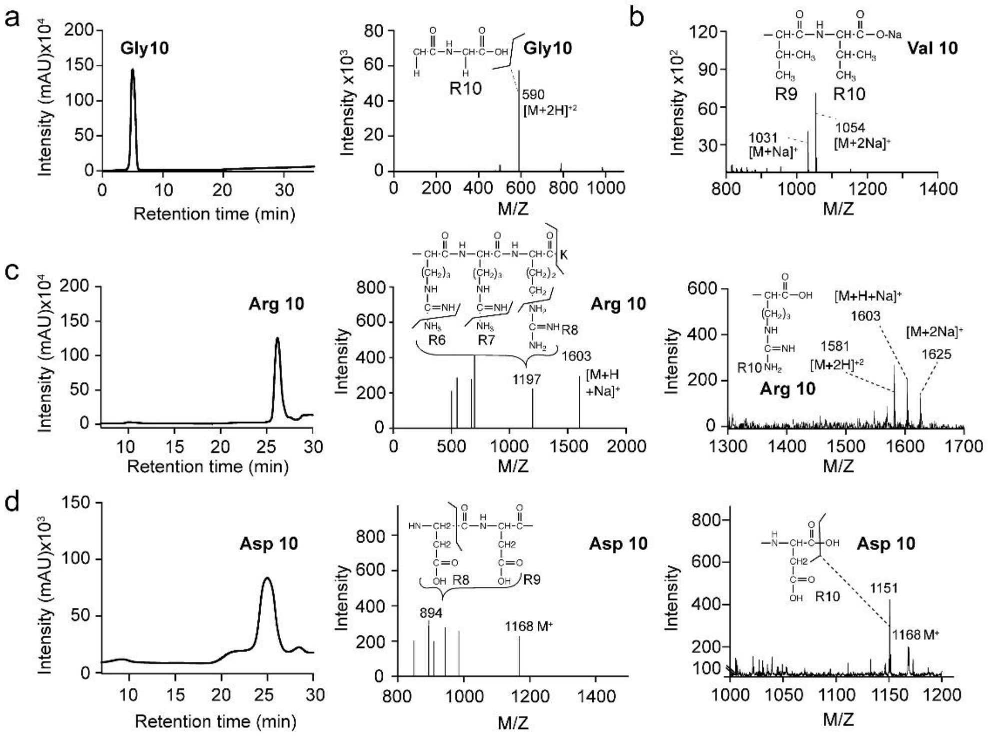


**Figure S14. Purity and characterization of polypeptides.** (a) HPLC chromatogram of $(Gly)_{10}$ and corresponding MALDI profile showing molecular weight peak at 590 ($MH^{+2}$). (b) MALDI profile of $(Val)_{10}$ showing molecular weight peak at 1031 ($MNa^{+}$). (c) HPLC chromatogram of $(Arg)_{10}$ and corresponding ESI (middle) and MALDI profile (right) showing molecular weight peak at 1603 ($MH^{+}Na^{+}$). (d) HPLC chromatogram of $(Asp)_{10}$ and corresponding ESI (middle) and MALDI profile (right) showing molecular weight peak at 1168 (M).

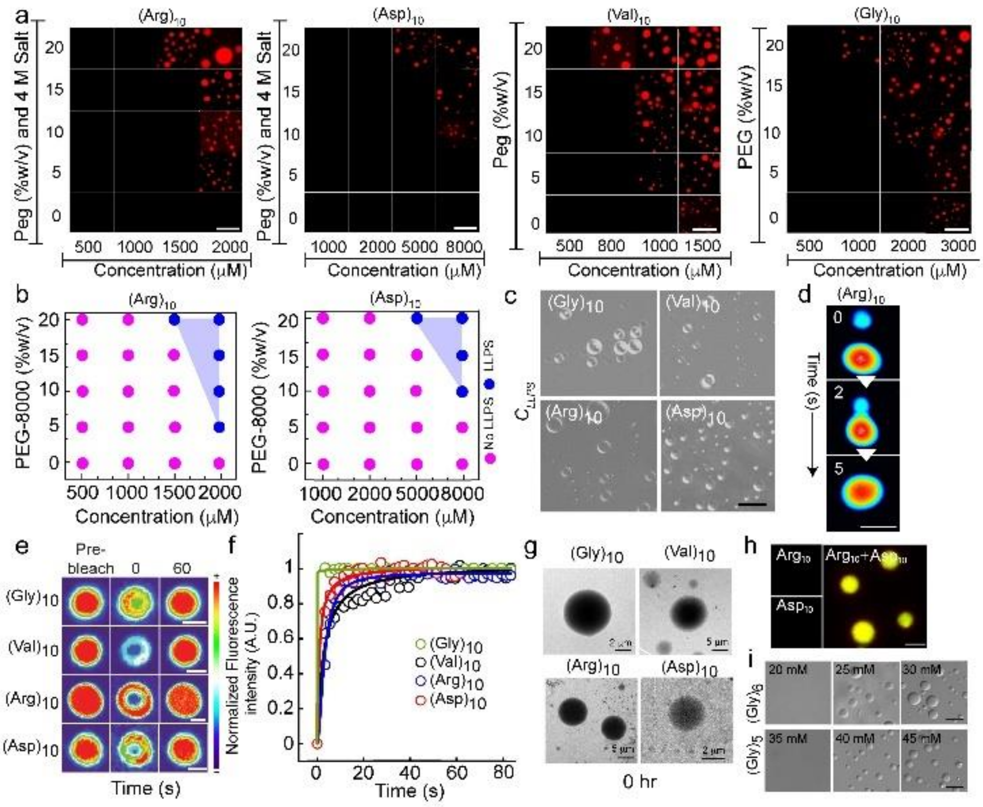


**Figure S15. LLPS of polypeptides *in vitro*** (a) Fluorescence microscopy images showing liquid condensate formation by the NHS-Rhodamine labeled [1:10 (v/v) labeled to unlabeled] polypeptides at varying PEG-8000 concentration (0%, 5%, 10%, 15% and 20% w/v). Notably, the phase separation of (Arg)$_{10}$ and (Asp)$_{10}$ was only observed in presence of 4 M NaCl. The scale bar is 5 μm. (b) Schematic showing phase regime of (Arg)$_{10}$ and (Asp)$_{10}$ in presence of 4 M NaCl. Different PEG-8000 and polypeptide concentrations were used for determining the phase regime. The pink color indicates no LLPS (soluble state) and the blue color indicates LLPS (condensate state). (c) DIC microscopy images of phase separated condensates of polypeptides at their respective $C_{LLPS}$ at 10% (w/v) PEG-8000 are shown. Notably, the phase separation condition for (Arg)$_{10}$ and (Asp)$_{10}$ was achieved only in presence of 4 M NaCl. The scale bar is 5 μm. *n=2*, independent experiments. (d) Time-lapse images of (Arg)$_{10}$ condensate depicting a fusion of condensate and subsequent growth over time. Represented in 'royal' LUT for better visualization. (e) Representative image showing the liquid nature of condensates at 48 h during FRAP analysis (before bleaching, at bleaching, and after bleaching). The images are represented in 'thermal' LUT for better visualization. The scale bar is 2 μm. (f) Normalized FRAP curve of phase separated condensates of polypeptides at 48 h. (g) Representative TEM images of the condensates formed by polypeptides immediately after LLPS (0 h) are shown. (h) Confocal microscopy showing co-LLPS of (Arg)$_{10}$ and (Asp)$_{10}$ [1:10 (v/v) labeled to unlabeled] at their respective $C_{LLPS}$ when mixed (in absence of salt) are shown. Respective polypeptide at $C_{LLPS}$ without mixing (in absence of salt) was used as a control. The scale bar is 5 µm. The experiment was repeated three times with similar observations. (i) DIC images of $(Gly)_5$ and $(Gly)_6$ at increasing polypeptide concentrations in presence of 10% (w/v) PEG-8000. The scale bar is 5 µm, $n=3$ independent experiments.

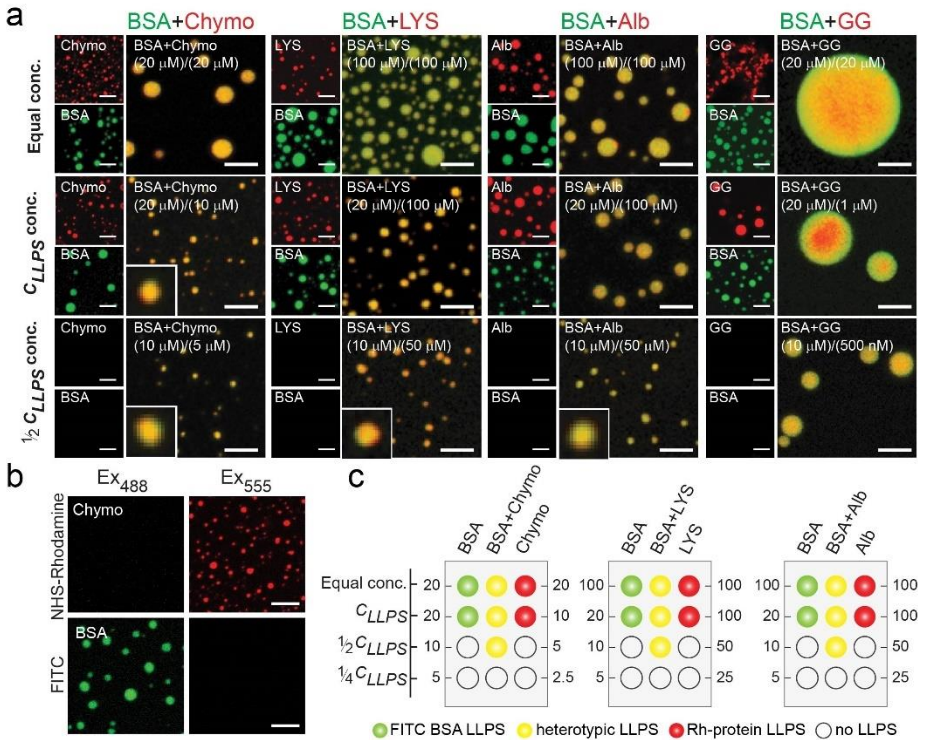

**Figure S16. Heterotypic co-LLPS by various combinations of proteins.** (a) Representative fluorescence microscopic images showing heterotypic co-LLPS by FITC labeled BSA and NHS-Rhodamine labeled various proteins [1:10 (v/v) labeled to unlabeled protein]. The confocal images showing co-LLPS of proteins when they were mixed at various protein concentrations (equal concentration, $C_{LLPS}$, and ½ $C_{LLPS}$) in 20 mM sodium phosphate buffer (pH 7.4) and 10% (w/v) PEG-8000. In BSA+GG co-LLPS, a red core-like structure was observed, which might be due to the preferential segregation of GG at the center of the heterotypic droplets. The scale bar is 5 µm. The experiment was repeated two times with similar observations. (b) Fluorescence microscopic images of the NHS-Rhodamine labeled Chymo (20 µm) and FITC labeled BSA (20 µm) at respective excitation wavelength and filter settings showing no bleed-through signal in the other channel during image acquisition. Similar settings were used during the acquisition of all the heterotypic condensates. The scale bar is 5 µm. (c) Schematic showing concentration requirement for LLPS in single protein component and two-protein component system. The color green indicates LLPS of FITC labeled BSA only (single component), red indicates LLPS of NHS-Rhodamine labeled protein (single component) and yellow indicates heterotypic co-LLPS. The empty symbol indicates no LLPS.

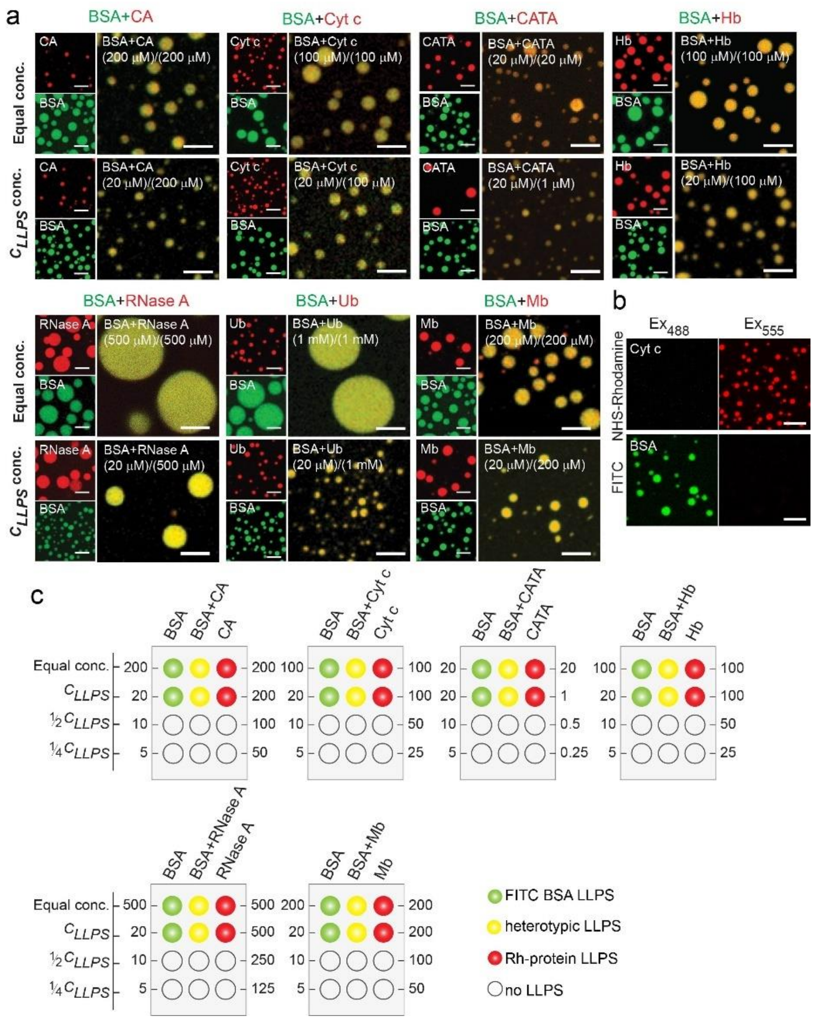

**Figure S17. Heterotypic co-LLPS by various combinations of proteins.** (a) Representative fluorescence microscopic images showing heterotypic phase separation by FITC labeled BSA and NHS-Rhodamine labeled various proteins [1:10 (v/v) labeled to unlabeled protein]. The confocal images showing co-LLPS of proteins when they were mixed at various protein concentrations (equal concentration and $C_{LLPS}$) in 20 mM sodium phosphate buffer (pH 7.4) and 10% (w/v) PEG-8000. The scale bar is 5 µm. The experiment was repeated two times with similar observations. (b) Fluorescence microscopic images of the NHS-Rhodamine labeled Cyt c (100 µm) and FITC labeled BSA (20 µm) at respective excitation wavelength and filter settings showing no bleed-through signal in the other channel during image acquisition. Similar settings were used during the acquisition of all the heterotypic condensates. The scale bar is 5 µm. (c) Schematic showing concentration requirement for LLPS in single protein component and two-protein component system. The color green indicates LLPS of FITC labeled BSA only (single component), red indicates LLPS of NHS-Rhodamine labeled protein (single component) and yellow indicates heterotypic co-LLPS. The empty symbol indicates no LLPS.

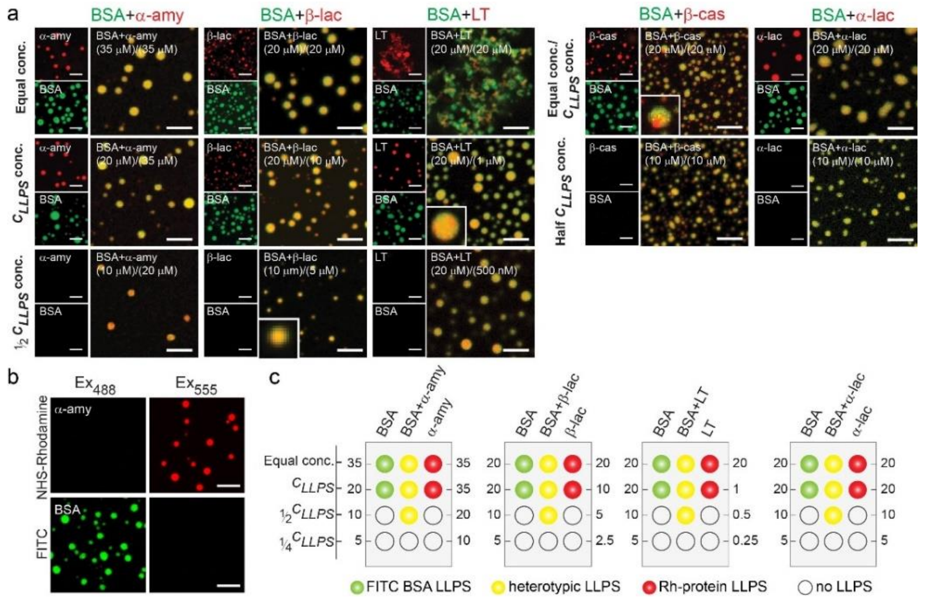


**Figure S18. Heterotypic co-LLPS by various combinations of proteins.** (a) Representative fluorescence microscopic images showing heterotypic co-LLPS phase separation by FITC labeled BSA and NHS-Rhodamine labeled various proteins [1:10 (v/v) labeled to unlabeled protein]. The confocal images showing co-LLPS of proteins when they were mixed at various protein concentrations (equal concentration, $C_{LLPS}$, and ½ $C_{LLPS}$) in 20 mM sodium phosphate buffer (pH 7.4) and 10% (w/v) PEG-8000. The scale bar is 5 µm. The experiment was repeated two times with similar observations. (b) Fluorescence microscopic images of the NHS-Rhodamine labeled α-amy (35 µm) and FITC labeled BSA (20 µm) at respective excitation wavelength and filter settings showing no bleed-through signal in the other channel during image acquisition. Similar settings were used during the acquisition of all the heterotypic condensates. The scale bar is 5 µm. (c) Schematic showing concentration requirement for LLPS in single protein component and two-protein component system. The color green indicates LLPS of FITC labeled BSA only (single component), red indicates LLPS of NHS-Rhodamine labeled protein (single component) and yellow indicates heterotypic co-LLPS. The empty symbol indicates no LLPS.

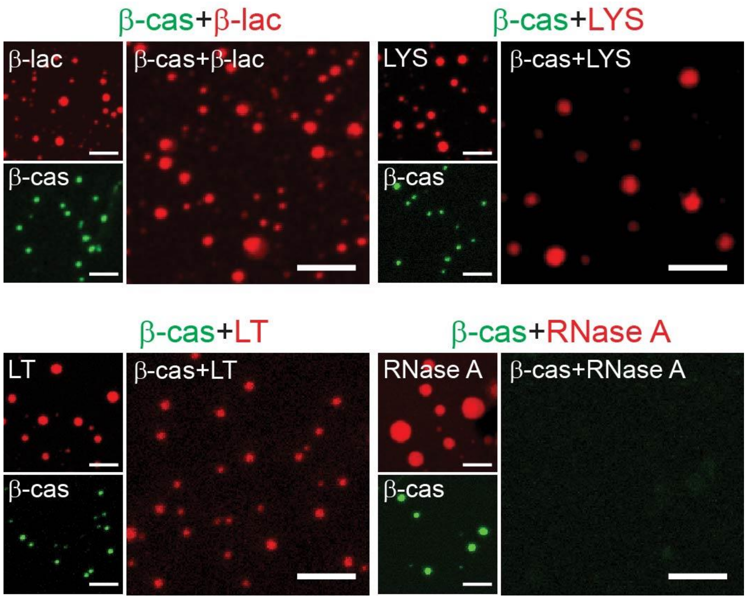


**Figure S19. Heterotypic co-LLPS by various combinations of proteins.** (a) Representative fluorescence microscopic images showing presence and absence of heterotypic co-LLPS phase separation by FITC labeled β-cas and NHS-Rhodamine labeled various proteins [1:10 (v/v) labeled to unlabeled protein]. Co-LLPS experiments were done with proteins mixed at their $C_{LLPS}$, in 20 mM sodium phosphate buffer (pH 7.4) and 10% (w/v) PEG-8000. The scale bar is 5 µm. The experiment was repeated two times with similar observations.

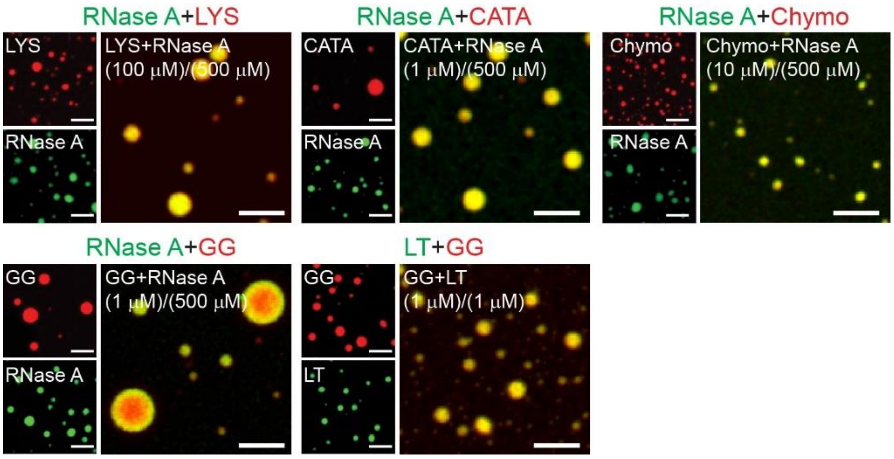


**Figure S20. Heterotypic co-LLPS by various combinations of proteins.** (a) Representative fluorescence microscopic images showing heterotypic co-LLPS phase separation by FITC labeled proteins (RNase A and LT) and NHS-Rhodamine labeled various proteins (LYS, CATA, Chymo and GG) [1:10 (v/v) labeled to unlabeled protein]. The confocal images showing co-LLPS of proteins when they were mixed at $C_{LLPS}$, in 20 mM sodium phosphate buffer (pH 7.4) and 10% (w/v) PEG-8000. The scale bar is 5 µm. The experiment was repeated two times with similar observations.

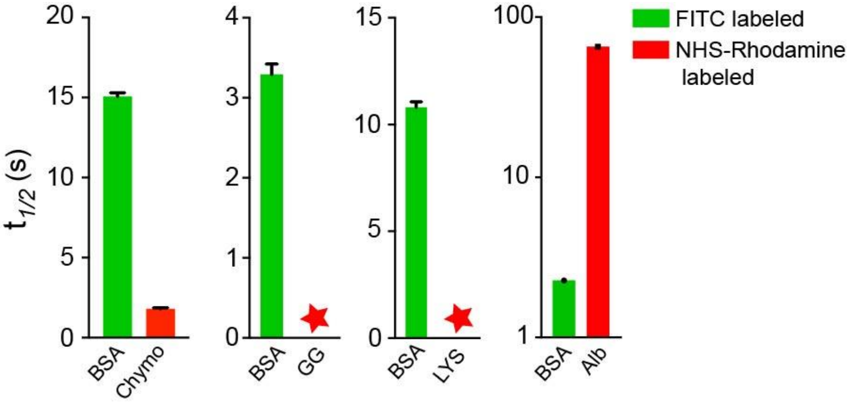


**Figure S21. Translational dynamics of selected proteins in the heterotypic liquid condensates.** The $t_{1/2}$ values were calculated from FRAP of heterotypic condensates. BSA and Alb showing slow fluorescence recovery in BSA+Chymo and BSA+Alb heterotypic condensates, respectively. Notably, $t_{1/2}$ values could not be calculated for GG, and LYS in the heterotypic condensate due to the negligible recovery after photobleaching. The data represent the mean ± *s.d.* for *n=2* independent experiments.

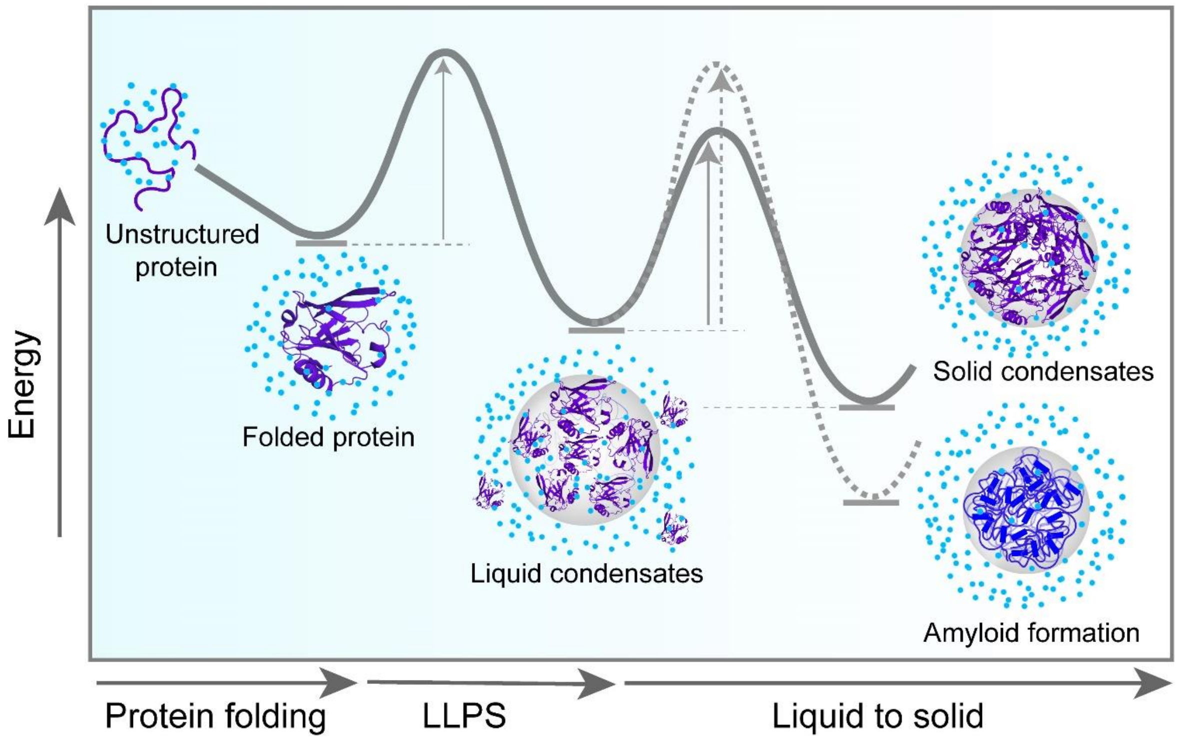


**Figure S22. Liquid-liquid phase separation and liquid-to-solid transition by proteins.** The schematic represents how folded proteins might undergo LLPS and liquid-to-solid transition. Blue dots represent water molecules (either surface-bound or free in solution). Solidification of condensates can result due to crystal packing or amorphous aggregation. Some proteins can also undergo solidification due to amyloid aggregation. The diagram indicates different protein states and the corresponding energy barrier for their interconversion.

**Table S1. Various proteins were used for the LLPS study showing their origin, molecular weight, and corresponding sequence sources.**

| S No | Protein | Catalog no. (Sigma) | Abbreviation | Origin | Residue | M.W. (Daltons) | Source (Uniprot ID) |
|---|---|---|---|---|---|---|---|
| 1 | Ribonuclease A | R6513 | RNase A | Bovine | 124 | 13690.29 | P61823 (27-150) |
| 2 | Lysozyme | L1667 | LYS | Human | 130 | 14700.67 | P61626 (19-148) |
| 3 | β-casein | C6905 | β-cas | Bovine | 209 | 23583.29 | P02666 (16-224) |
| 4 | Lactoferrin | L4040 | LT | Human | 710 | 78181.95 | P02788 (1-710) |
| 5 | BSA | TC194 (Himedia, India) | BSA | Bovine | 583 | 66432.96 | P02769 (25-607) |
| 6 | Catalase | C40 | CATA | Bovine | 506 | 57583.66 | P00432 (2-527) |
| 7 | β-lactoglobulin | L3908 | β-lac | Bovine | 162 | 18281.21 | P02754 (17-178) |
| 8 | α-lactalbumin | L5385C2506 | α-lac | Bovine | 123 | 14186.06 | P00711 (20-142) |
| 9 | Myoglobin | M1882 | Mb | Equine heart | 153 | 16951.48 | P68082 (2-154) |
| 10 | Hemoglobin | H7379 | Hb | Human | 141 (α chain)<br>146 (β-chain) | 15126.36 (α chain)<br>15867.22 (β-chain) | P69905 (α-chain) (2-147)<br>P68871 (β-chain) (2-142) |
| 11 | α-amylase | A4551 | α-amy | *Bacillus lichenoformis* | 483 | 55268.17 | P06278 (30-512) |
| 12 | Chymotrypsin | SKU1023070001 | Chymo | Bovine pancreas | 241 | 25207.64 | P00766 (chain A-1-13, chain B-16-146, chain C-149-245) |
| 13 | α-Synuclein | Expressed and purified | α-Syn | Human | 140 | 14460.16 | P37840 (1-140) |

| | | | | | | | |
|---|---|---|---|---|---|---|---|
| 14 | Tau-F | Expressed and purified | Tau | Human | 441 | 45849.91 | P10636-8 (1-441) |
| 15 | Ovalbumin | A5503 | Alb | Chicken | 385 | 42750.04 | P01012 (2-386) |
| 16 | Ubiquitin | U6253 | Ub | Bovine erythrocytes | 76 | 8530.83 | A0A3Q1 M4K3 (1-76) |
| 17 | Cytochrome c | C2506 | Cyt c | Equine | 104 | 11701.55 | P00004 (2-105) |
| 18 | γ-globulin | G5009 | GG | Bovine | * | 150000 | * |
| 19 | Carbonic anhydrase | C3934 | CA | Bovine | * | 30000 | * |

*Notably, sequence information was not available for γ-globulin and carbonic anhydrase.

**Table S2. Structural and biophysical characterization of the proteins.** The secondary structure of the proteins was determined by CD spectroscopy. The primary sequence analysis of the proteins was done using IUPred2A[17] to predict the disorder tendency; SMART[18] to analyze the presence of low complexity domains (LCDs) and CatGranule[19] for predicting the propensity for LLPS.

| S. No. | Protein | Secondary structure | PI | Positive charge (P) | Negative charge (N) | LCD | IDR | CatGranule Score | $\sqrt{(N^2+P^2)}$ |
|---|---|---|---|---|---|---|---|---|---|
| 1 | RNase A | Helix | 8.64 | 14 | 10 | ✓ | ✓ | -0.93844 | 17.20 |
| 2 | LYS | Helix | 9.28 | 19 | 11 | ✕ | ✕ | -0.0412797 | 21.95 |
| 3 | β-cas | Random coil | 5.13 | 15 | 23 | ✓ | ✓ | -1.15637 | 27.45 |
| 4 | LT | Helix | 8.5 | 90 | 79 | ✓ | ✓ | 0.827097 | 119.75 |
| 5 | BSA | Helix | 5.6 | 82 | 99 | ✓ | ✕ | -0.273782 | 128.54 |
| 6 | CATA | Helix | 6.63 | 58 | 62 | ✕ | ✓ | 0.960901 | 84.89 |
| 7 | β-lac | Helix | 4.83 | 18 | 26 | ✕ | ✕ | -1.22001 | 31.62 |
| 8 | α-lac | Helix | 4.8 | 13 | 20 | ✓ | ✕ | -0.441139 | 29.69 |
| 9 | Mb | Helix | 7.36 | 21 | 21 | ✕ | ✓ | 0.551681 | 0 |
| 10 | Hb | Helix | 8.13 | 56 | 54 | ✕ | ✕ | 0.48402 | 77.79 |
| 11 | α-amy | Helix | 6.05 | 50 | 62 | ✕ | ✓ | 1.45616 | 79.64 |
| 12 | Chymo | Random coil | 8.33 | 17 | 14 | ✕ | ✓ | 0.151479 | 22.02 |
| 13 | α-Syn | Random coil | 4.67 | 15 | 24 | ✓ | ✓ | 1.12517 | 28.30 |
| 14 | Tau | Random coil | 8.24 | 58 | 56 | ✓ | ✓ | 1.68922 | 80.62 |
| 15 | Alb | Helix | 5.19 | 35 | 47 | ✓ | ✓ | -0.234022 | 58.60 |
| 16 | Ub | Helix | 6.56 | 11 | 11 | ✕ | ✓ | -0.472928 | 15.55 |
| 17 | Cyt c | Helix | 9.59 | 21 | 12 | ✕ | ✓ | 1.15134 | 24.18 |

**Table S3. Heterotypic co-LLPS by various combinations of proteins. The symbol (✓) and (×) indicates presence and absence of condensate, respectively. The experiment was done with proteins at their respective $C_{LLPS}$.**

| Protein partner (1) (FITC) | Protein partner (2) (NHS-Rh) | Heterotypic co-LLPS |
|---|---|---|
| β-cas (✓) | β-lac (✓) | × (Only red condensates) |
| β-cas (✓) | LT (✓) | × (Only red condensates) |
| β-cas (✓) | LYS (✓) | × (Only red condensates) |
| β-cas (✓) | RNase A (✓) | × (No condensates) |
| LT (✓) | GG (✓) | × (Yellow condensates) |
| RNase A (✓) | LYS (✓) | ✓ (Yellow condensates) |
| RNase A (✓) | CATA (✓) | ✓ (Yellow condensates) |
| RNase A (✓) | Chymo (✓) | ✓ (Yellow condensates) |
| RNase A (✓) | GG (✓) | ✓ (Condensates with red core) |
